# The IRE1/XBP1 signaling axis promotes skeletal muscle regeneration through a cell non-autonomous mechanism

**DOI:** 10.1101/2021.08.19.457023

**Authors:** Anirban Roy, Meiricris Tomaz da Silva, Raksha Bhat, Kyle R. Bohnert, Takao Iwawaki, Ashok Kumar

## Abstract

Skeletal muscle regeneration is regulated by coordinated activation of multiple signaling pathways activated in both injured myofibers and satellite cells. The unfolded protein response (UPR) is a major mechanism that detects and alleviates protein-folding stresses in ER. However, the role of individual arms of the UPR in skeletal muscle regeneration remains less understood. In the present study, we demonstrate that IRE1α (also known as ERN1) and its downstream target, XBP1, are activated in skeletal muscle of mice upon injury. Myofiber-specific ablation of IRE1 or XBP1 in mice diminishes skeletal muscle regeneration that is accompanied with reduced number of satellite cells and their fusion to injured myofibers. Ex vivo cultures of myofiber explants demonstrate that ablation of IRE1α reduces the proliferative capacity of myofiber-associated satellite cells. Myofiber-specific deletion of IRE1α dampens Notch signaling and canonical NF-κB pathway in skeletal muscle of mice. Our results also demonstrate that targeted ablation of IRE1α reduces skeletal muscle regeneration in the mdx mice, a model of Duchenne muscular dystrophy. Collectively, our results reveal that the IRE1α-mediated signaling promotes muscle regeneration through augmenting the proliferation of satellite cells in a cell non-autonomous manner.

## Introduction

Skeletal muscle, the most abundant tissue of the body, has remarkable regenerative capacity mainly due to its resident muscle stem cells, also known as satellite cells. These cells are located beneath the basal lamina of myofiber in a dormant state. Upon muscle injury, satellite cells become activated leading to their asymmetric cell division and differentiation into myoblasts which eventually fuse together or with existing myofibers to accomplish muscle regeneration (Kuang and Rudnicki, 2008; Yin et al., 2013). In many muscle diseases, such as Duchenne muscular dystrophy (DMD) and aged skeletal muscle (sarcopenia), the abundance and regenerative capacity of satellite cells is diminished leading to impairment in muscle regeneration and a net loss of muscle mass (Brack and Munoz-Canoves, 2016; Dumont et al., 2015). It is now increasingly evidenced that muscle regeneration is regulated through orchestrated activation of multiple signaling pathways and factors produced by regenerating myofiber as well as other cell types (Dumont et al., 2015). However, the mechanisms of muscle regeneration are not yet fully understood.

The endoplasmic reticulum (ER) is the major subcellular compartment that is involved in protein folding and secretion. Many secreted proteins are translated from mRNAs localized on the cytosolic side of the ER membrane and enter the ER as nascent chains that are modified and properly folded before exiting the organelle (Harding et al., 1999). An imbalance between the influx of unfolded proteins and the capacity of the organelle to handle them cause stress in the ER that triggers activation of an adaptive signaling network, known as the unfolded protein response (UPR) (Hetz, 2012; Preissler and Ron, 2019; Wang and Kaufman, 2014). The major goal of the UPR is to restore homeostasis or induce apoptosis of permanently damaged cells (Tabas and Ron, 2011). The UPR is initiated by three ER transmembrane sensors: RNA-dependent protein kinase-like ER eukaryotic translation initiation factor 2 alpha kinase (PERK), inositol-requiring enzyme 1 (IRE1), and activating transcription factor 6 (ATF6). PERK directly phosphorylates eukaryotic translation initiation factor 2 α (eIF2α), which leads to the attenuation of translation initiation and limits the protein-folding load on the ER. IRE1α (also known as ERN1, Endoplasmic Reticulum to Nucleus signaling 1) is a serine/threonine kinase and endoribonuclease controlling cell fate under ER stress. Once activated, IRE1α oligomerizes leading to three major downstream outputs: the activation of c-Jun N-terminal kinase (JNK), the splicing of *Xbp1* mRNA, and the degradation of targeted mRNA and microRNAs, a process referred to as regulated IRE1-dependent decay (RIDD) (Hollien et al., 2009; Hollien and Weissman, 2006; Maurel et al., 2014; Wang and Kaufman, 2014). Upon stress in ER, ATF6 protein is transported to the Golgi complex where it is cleaved by S1P and S2P to release the N-terminal-domain. The cleaved ATF6 domain translocates to the nucleus, where in association with other factors, it induces gene transcription of target molecules such as GRP78, GRP94 and calnexin (Hetz, 2012; Walter and Ron, 2011; Wang and Kaufman, 2014). Although ER stress is the major stimuli, a few components of the UPR, including IRE1α and XBP1 can also be activated in the absence of ER stress (Abdullah and Ravanan, 2018; Dufey et al., 2020; Martinon et al., 2010).

Because regeneration of injured skeletal muscle involves a huge increase in demand for synthesis, processing, and secretion of multiple growth factors and signaling proteins and synthesis of an entirely new set of contractile, cytoskeletal, and membrane proteins, the activation of UPR may be a physiological response to ensure that cells participating in regeneration continue to function efficiently during these increased demands. Indeed, accumulating evidence suggests that the UPR is activated and plays an important role in the regulation of skeletal muscle mass and myogenesis (Afroze and Kumar, 2019; Bohnert et al., 2018). ATF6 arm of the UPR is activated during skeletal muscle development where it mediates apoptosis of a subpopulation of myoblasts that may be incompetent of handling cellular stresses (Nakanishi et al., 2005). The role of the UPR in myogenesis is also supported by the findings that pan-inhibition of ER stress attenuates apoptosis and myogenic differentiation (Nakanishi et al., 2005) whereas inducers of ER stress remove vulnerable myoblasts allowing more efficient differentiation into myotubes in cell cultures (Nakanishi et al., 2007). Recent studies have suggested that PERK arm of the UPR is required for maintaining satellite cells in a quiescent state in skeletal muscle of adult mice. Genetic inducible ablation of PERK in satellite cells inhibits skeletal muscle regeneration in adult mice (Xiong et al., 2017; Zismanov et al., 2016).

Moreover, PERK-ATF4 axis regulates the early differentiation of C2C12 myoblasts potentially through up-regulating the expression of differentiation-associated microRNAs (Tan et al., 2021). Intriguingly, a very recent study demonstrated that IRE1α promotes skeletal muscle regeneration through RIDD-mediated reduction in myostatin levels (He et al., 2021). However, the role and the mechanisms by which the IRE1α/XBP1 signaling in myofiber regulates skeletal muscle regeneration in adult mice remain poorly understood.

In the present study, using genetic mouse models, we have investigated the role of IRE1 and XBP1 in skeletal muscle regeneration. Our results demonstrate myofiber-specific ablation of IRE1α attenuates skeletal muscle regeneration in adult mice. IRE1α controls muscle regeneration through the activation of XBP1 transcription factor, but not through RIDD mechanism. Genetic ablation of *Xbp1* also reduces regeneration of injured skeletal muscle. Our results also demonstrate that IRE1α/XBP1 pathway in myofiber regulates the proliferation of satellite cells in a non-cell-autonomous manner. Deletion of IRE1α in myofibers suppresses Notch and canonical NF-κB signaling in regenerating skeletal muscle. Finally, our experiments demonstrate that genetic ablation of IRE1α exacerbates myopathy and reduces the number of satellite cells in skeletal muscle of dystrophin-deficient mdx (a model of DMD) mice.

## Results

### Myofiber-specific deletion of IRE1α attenuates skeletal muscle regeneration in adult mice

We first investigated how the phosphorylation of IRE1α and levels of spliced XBP1 (sXBP1) protein are regulated in skeletal muscle after injury. Left side tibialis anterior (TA) muscle of wild-type (WT) mice was injected with 1.2% BaCl_2_ solution, a widely used myotoxin for experimental muscle injury (Hindi et al., 2017; Xiong et al., 2017), whereas contralateral TA muscle was injected with saline only to serve as control. Results showed that the levels of phosphorylated IRE1α were significantly increased in TA muscle at day 5 post-injury compared to contralateral control muscle. Furthermore, the levels of spliced XBP1 (sXBP1) protein were also found to be considerably increased in injured TA muscle of mice suggesting activation of the IRE1α/XBP1 pathway in injured muscle (**Figure 1A**).

**FIGURE 1.**
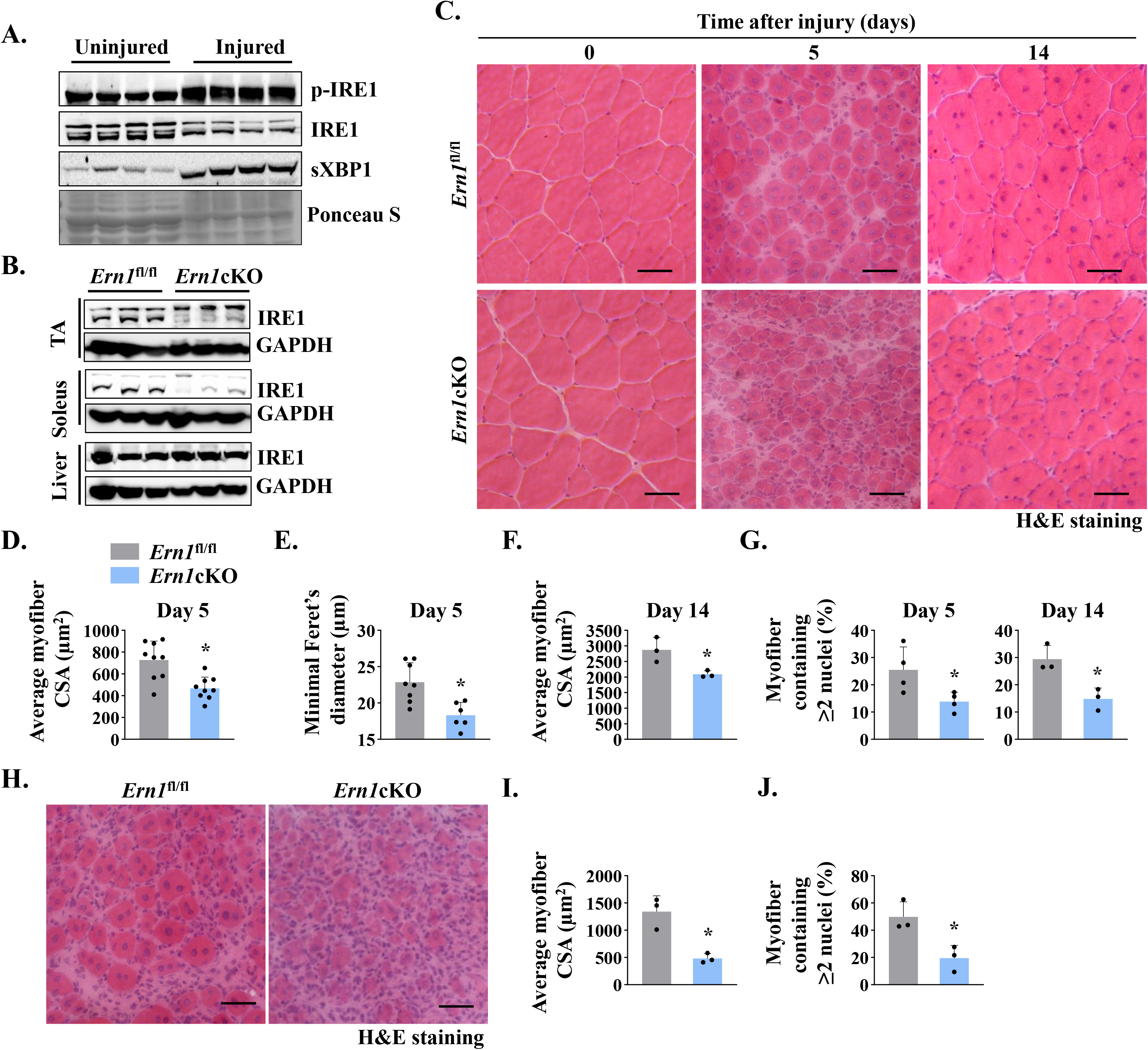
IRE1α is required for muscle regeneration. (**A**) Left side TA muscle of wild-type mice was injected with 50 µl of 1.2% BaCl_2_ solution whereas contralateral TA muscle was injected with normal saline to serve as uninjured control. After 5d, the TA muscles were harvested and analyzed by performing western blotting. Representative immunoblots presented here demonstrate the levels of p-IRE1, IRE1, and sXBP1 protein in uninjured and injured TA muscle. Ponceau S staining confirmed equal loading of protein in each lane (n=4 mice per group). (**B**) Immunoblots presented here show levels of IRE1α protein in TA and soleus muscle and liver of *Ern1*^fl/fl^ and *Ern1*cKO mice. GAPDH was used as loading control (n=3 mice per group). (**C**) Representative photomicrographs of Hematoxylin and Eosin (HE)-stained sections of TA muscles of *Ern1*^fl/fl^ and *Ern1*cKO mice at day 0, 5, and 14 after intramuscular injection of BaCl_2_ solution. Scale bar: 50 µm. Quantification of average myofiber (**D**) cross-sectional area (CSA) (n=9 mice per group) and (**E**) Minimal Feret’s diameter in TA muscle at day 5 post-injury (n=6 mice per group). (**F**) Average myofiber CSA in TA muscle of *Ern1*^fl/fl^ and *Ern1*cKO mice after 14 days of BaCl_2_-mediated injury (n=3 mice per group). (**G**) Percentage of myofibers containing 2 or more centrally located nuclei in TA muscle at day 5 (n=4 mice per group) and day 14 (n=3 mice per group) after BaCl_2_-mediated injury. After 21 days of first injury, TA muscle of *Ern1*^fl/fl^ and *Ern1*cKO mice was again given intramuscular injection of 50 µl of 1.2% BaCl_2_ solution, and the muscle was analyzed at day 5. Representative photomicrograph of (**H**) H&E-stained TA muscle sections and quantitative estimation of **(I)** average myofiber CSA and **(J)** percentage of myofibers containing two or more centrally located nuclei. Scale bar: 50 µm. n=3 mice per group. Data are presented as mean ± SEM. *p≤0.05, values significantly different from corresponding injured TA muscle of *Ern1*^fl/fl^ mice by unpaired t test.

We next sought to investigate the role of IRE1α (gene name: *Ern1*) in skeletal muscle regeneration. Floxed *Ern1* (henceforth *Ern1*^fl/fl^) mice were crossed with muscle creatine kinase (*Mck*)-Cre to generate *Ern1*^fl/fl^;*Mck*-Cre (henceforth *Ern1*cKO) and littermate *Ern1*^fl/fl^ mice. There was no significant difference in their body weight or muscle weight of *Ern1*^fl/fl^ and *Ern1*cKO mice in naïve conditions (**Figure 1-figure supplement 1A, 1B**). Western blot analysis confirmed that levels of IRE1α protein were considerably reduced in TA and soleus muscle, but not in liver of *Ern1*cKO mice (**Figure 1B**). To understand the role of IRE1α signaling in muscle regeneration, TA muscle of 3-month old littermate *Ern1*^fl/fl^ and *Ern1*cKO mice was injected with 50 µl of 1.2% BaCl_2_ solution to induce necrotic muscle injury. Muscle regeneration was evaluated at day 5 and day 14 post-BaCl_2_ injection by performing Hematoxylin and Eosin (H&E) staining (**Figure 1C**). Results showed that the regeneration of TA muscle was attenuated in *Ern1*cKO mice compared to *Ern1*^fl/fl^ mice at day 5 post-injury. There was an apparent decrease in the size of centronucleated myofibers (CNFs) and an increase in the cellular infiltrate in *Ern1*cKO mice compared to *Ern1*^fl/fl^ mice (**Figure 1C**). Morphometric analysis H&E-stained TA muscle sections revealed a significant decrease in the average cross-sectional area (CSA) and minimal Feret’s diameter of regenerating myofibers in *Ern1*cKO mice compared with *Ern1*^fl/fl^ mice (**Figure 1D-F**). Moreover, the percentage of myofibers containing two or more centrally located nuclei was significantly reduced in injured TA muscle of *Ern1*cKO mice compared with *Ern1*^fl/fl^ mice (**Figure 1G**). Deficits in muscle regeneration in *Ern1*cKO mice were also present at day 14 after muscle injury (**Figure 1C, 1F & 1G**).

Successive muscle injuries are used to study satellite cell pool maintenance or depletion after more than one round of regeneration. In this approach, second injury is carried out three to four weeks apart, which is enough time elapse to allow the regeneration of the muscle after first injury (Hardy et al., 2016; Hindi and Kumar, 2016). Therefore, we next examined muscle regeneration in *Ern1*^fl/fl^ and *Ern1*cKO mice after performing double injury. At day 21 of first injury, TA muscle of *Ern1*^fl/fl^ and *Ern1*cKO mice was injured again by intramuscular injection of 50 μl 1.2% BaCl_2_ solution. After 5 days, the muscle was isolated and analyzed by performing H&E staining (**Figure 1H**). Results showed that defect in muscle regeneration in *Ern1*cKO mice was more pronounced after second round of injury (**Figure 1H-1J**). Moreover, Masson’s Trichrome staining of TA muscle sections showed considerably increase in deposition of fibrotic tissue at day 5 after second injury (**Figure 1-figure supplement 1C**).

### Ablation of IRE1α reduces the expression of early markers of muscle regeneration in mice

To further understand the role of IRE1α in skeletal muscle regeneration, we studied the expression of early markers of regeneration such embryonic myosin heavy chain (*Myh3*) and various myogenic regulatory factors (MRFs). We first analyzed 5d-injured TA muscle of *Ern1*^fl/fl^ and *Ern1*cKO mice by immunostaining for embryonic myosin heavy chain (eMyHC) protein and laminin protein. Nuclei were counterstained with DAPI (4′,6-diamidino-2-phenylindole). Results showed that the proportion of eMyHC^+^ myofibers within laminin staining with higher cross-sectional area (CSA) were reduced in TA muscle of *Ern1*cKO mice compared to *Ern1*^fl/fl^ mice (**Figure 2A, 2B**). Moreover, the percentage of eMyHC^+^ myofibers containing 2 or more centrally located nuclei was significantly reduced in regenerating TA muscle of *Ern1*cKO mice compared with *Ern1*^fl/fl^ mice (**Figure 2C**). Skeletal muscle regeneration involves a sequential up-regulation of *Myf5*, *Myod1*, *Myog* (myogenin), and *eMyHC*. Our qPCR analysis showed that there was no significant difference in the mRNA levels of *Myh3*, *Myf5*, *Myod1*, and *Myog* in uninjured TA muscle of *Ern1*^fl/fl^ and *Ern1*cKO mice. However, mRNA levels of *Myf5*, *Myod1*, *Myog*, and *Myh3* were found to be significantly reduced in 5d-injured TA muscle of *Ern1*cKO mice compared to 5d-injured TA muscle of *Ern1*^fl/fl^ mice (**Figure 2D-2G**). Furthermore, western blot analysis showed that protein levels of MYOD and eMyHC, but not myogenin, were considerably reduced in 5d-injured TA muscle of *Ern1*cKO mice compared to *Ern1*^fl/fl^ mice (**Figure 2H**). Collectively, these results suggest that myofiber-specific ablation of IRE1α attenuates regenerative myogenesis in adult mice.

**FIGURE 2.**
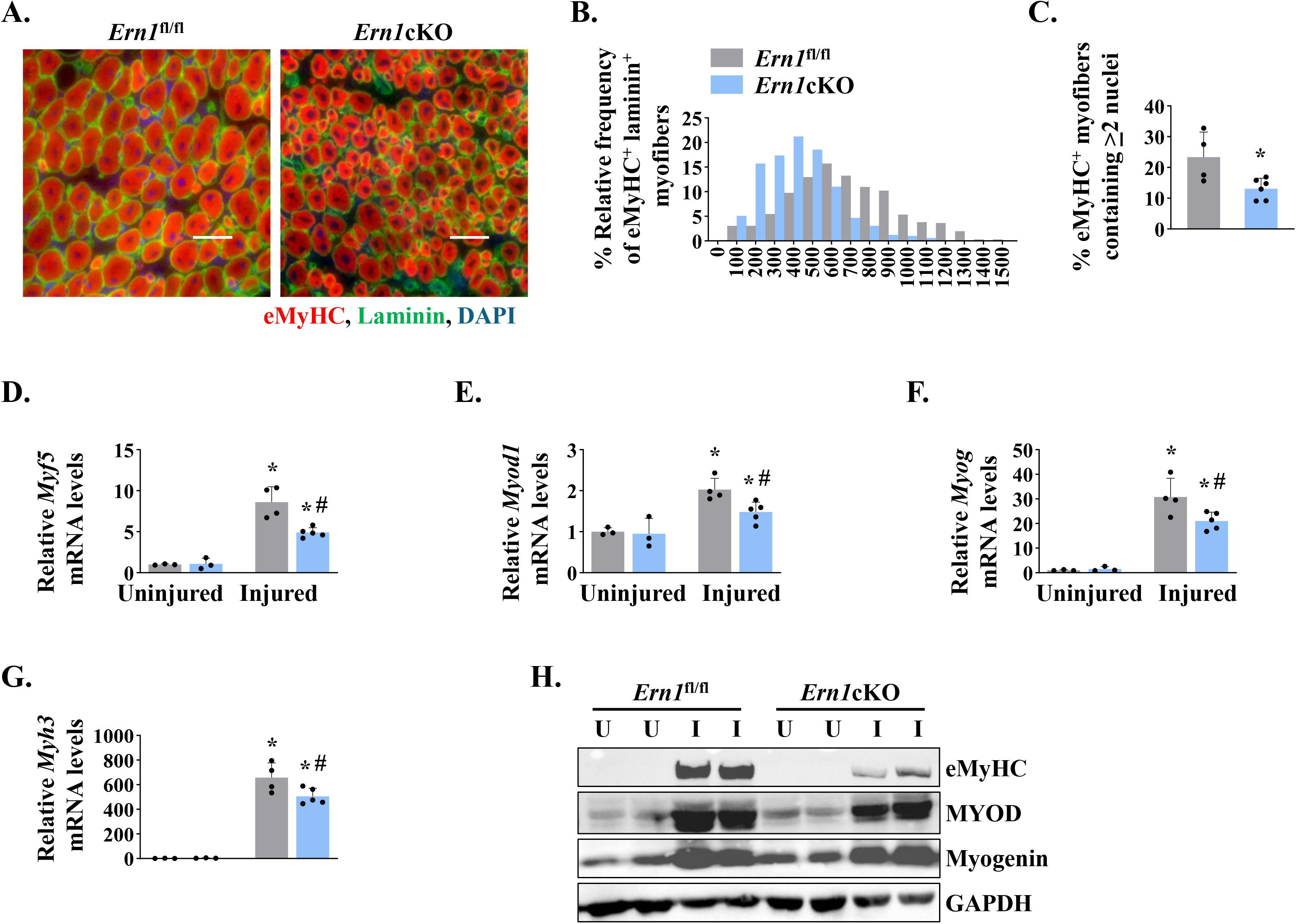
Myofiber-specific ablation of IRE1α inhibits early markers of muscle regeneration. (**A**) Representative photomicrographs of 5d-injured TA muscle sections of *Ern1*^fl/fl^ and *Ern1*cKO mice after immunostaining for eMyHC (red) and laminin (green) protein. Nuclei were counterstained with DAPI (blue). (**B**) Frequency distribution of eMyHC^+^ Laminin^+^ myofiber CSA in 5d-injured TA muscle of *Ern1*cKO and *Ern1*^fl/fl^ mice (n=4 for *Ern1*^fl/fl^ and n=3 for *Ern1*cKO group). (**C**) Percentage of eMyHC^+^ myofibers with 2 or more nuclei in 5d-injured TA muscle of *Ern1*^fl/fl^ and *Ern1*cKO mice (n=4 for *Ern1*^fl/fl^ and n=6 for *Ern1*cKO group). Data expressed as mean ± SEM. *P≤0.05, values significantly different from 5d-injured TA muscle of *Ern1*^fl/fl^ mice by unpaired t test. Relative mRNA levels of (**D**) *Myf5*, (**E**) *Myod1*, (**F**) *Myog*, and (**G**) *Myh3* in uninjured and 5d-injured TA muscle of *Ern1*^fl/fl^ and *Ern1*cKO mice. (**H**) Representative western blot showing levels of eMyHC, MyoD, myogenin, and GAPDH in uninjured and 5d-injured TA muscle of *Ern1*^fl/fl^ and *Ern1*cKO mice (n=3 for Uninjured *Ern1*^fl/fl^, n=3 for Uninjured *Ern1*cKO, n=4 for Injured *Ern1*^fl/fl^ and n=5 for Injured *Ern1*cKO group). Data are presented as mean ± SEM and analyzed by one-way analysis of variance (ANOVA) followed by Tukey’s multiple comparison test. **\***P≤0.05, values significantly different from uninjured TA muscle of *Ern1*^fl/fl^ mice. ^#^P ≤ 0.05, values significantly different from 5d-injured TA muscle of *Ern1*^fl/fl^ mice. U, uninjured; I, injured.

### Ablation of IRE1α reduces abundance of satellite cells in regenerating skeletal muscle

Since satellite cells are indispensable for muscle repair (Yin et al., 2013), we next investigated whether myofiber-specific ablation of IRE1α affects the number of satellite cells in injured skeletal muscle. Pax7 is a transcription factor that is expressed in both quiescent and activated satellite cells (Dumont et al., 2015). Indeed, Pax7 has been widely used as a marker to label and quantify satellite cells on muscle sections (Hindi and Kumar, 2016; Ogura et al., 2015).

Transverse sections generated from uninjured and 5d-injured TA muscle of *Ern1*^fl/fl^ and *Ern1*cKO mice were immunostained for Pax7 to detect satellite cells. In addition, the sections were immunostained for laminin to mark the boundary of the myofibers and DAPI was used to stain nuclei (**Figure 3A**). There was no significant difference in the number of satellite cells in uninjured TA muscle of *Ern1*^fl/fl^ and *Ern1*cKO mice (**Figure 3A**). However, the number of satellite cells per unit area was significantly reduced in injured TA muscle of *Ern1*cKO mice compared with injured TA muscle of *Ern1*^fl/fl^ mice (**Figure 3A, 3B**). Moreover, mRNA levels of *Pax7* were found to be significantly reduced in injured TA muscle of *Ern1*cKO mice compared to that of *Ern1*^fl/fl^ mice (**Figure 3C**). Similarly, protein levels of Pax7 were also markedly lower in 5d-injured TA muscle of *Ern1*cKO mice compared with *Ern1*^fl/fl^ mice (**Figure 3D**).

**FIGURE 3.**
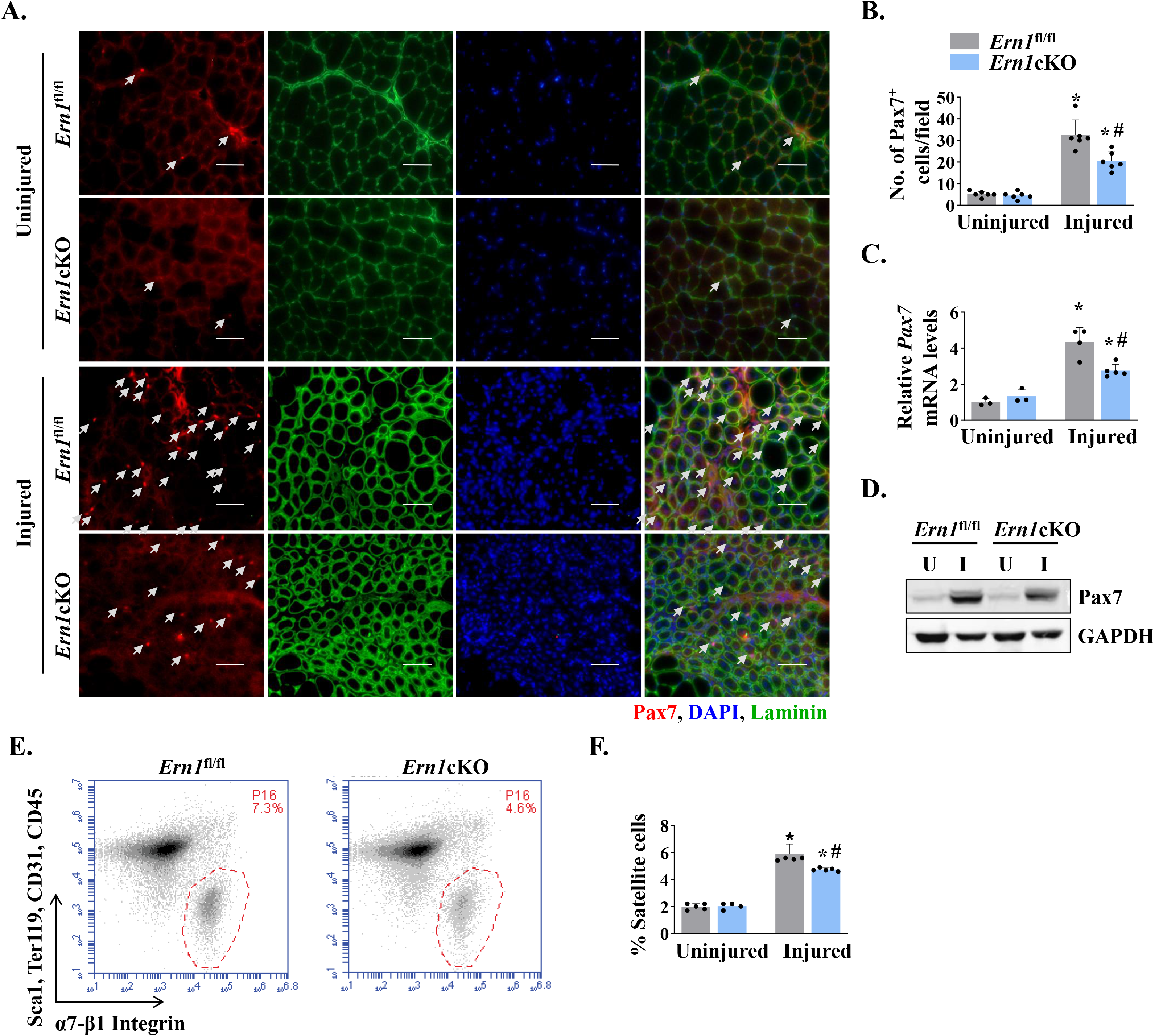
Myofiber-specific ablation of IRE1α reduces abundance of satellite cells in injured muscle. (**A**) Representative photomicrographs of uninjured and 5d-injured TA muscle sections of *Ern1*^fl/fl^ and *Ern1*cKO mice after immunostaining for Pax7 (red) and laminin (green) protein. Nuclei were identified by staining with DAPI. Scale bar: 50 µm. White arrows point to Pax7^+^ satellite cells. (**B**) Average number of Pax7^+^ cells per field (∼0.15 mm^2^) in uninjured and 5d-injured TA muscle of *Ern1*^fl/fl^ and *Ern1*cKO mice. (**C**) Relative mRNA levels of *Pax7* in uninjured and 5d-injured TA muscle of *Ern1*^fl/fl^ and *Ern1*cKO mice assayed by performing qPCR (n=3 for Uninjured *Ern1*^fl/fl^, n=3 for Uninjured *Ern1*cKO, n=4 for Injured *Ern1*^fl/fl^ and n=5 for Injured *Ern1*cKO group). (**D**) Representative immunoblots showing levels of Pax7 and unrelated protein GAPDH in uninjured and 5d-injured TA muscle of *Ern1*^fl/fl^ and *Ern1*cKO mice. (**E**) Primary mononuclear cells were isolated from TA muscle of *Ern1*^fl/fl^ and *Ern1*cKO mice and subjected to FACS analysis for satellite cells. Representative FACS dot plots demonstrating the percentage of α7-integrin^+^ cells in 5d-injured TA muscle of *Ern1*^fl/fl^ and *Ern1*cKO mice. (**F**) Quantification of α7-integrin^+^ satellite cells in uninjured and 5d-injured TA muscle of *Ern1*^fl/fl^ and *Ern1*cKO mice assayed by FACS (n=4 in each group). Data are expressed as mean ± SEM and analyzed by one-way analysis of variance (ANOVA) followed by Tukey’s multiple comparison test. **\***P≤0.05, values significantly different from uninjured TA muscle of *Ern1*^fl/fl^ mice. ^#^P ≤ 0.05, values significantly different from 5d-injured TA muscle of *Ern1*^fl/fl^ mice. U, uninjured; I, injured.

Fluorescence-activated cell sorting (FACS) analysis using a combination of cell surface markers (i.e. CD45^-^, CD31^-^, Ter119^−^, Sca-1^-^, and α7-β1 Integrin^+^) is another approach to quantify the number of satellite cells in skeletal muscle in mice (Hindi and Kumar, 2016; Ogura et al., 2015). We next performed FACS to quantify the number of satellite cells in skeletal muscle of *Ern1*^fl/fl^ and *Ern1*cKO mice. There was no significant difference in the numbers of satellite cells between uninjured TA muscle of *Ern1*^fl/fl^ and *Ern1*cKO mice (data not shown).

However, our analysis showed that proportion of satellite cells was significantly reduced in the 5d-injured TA muscle of *Ern1*cKO mice compared to injured TA muscle of *Ern1*^fl/fl^ mice (**Figure 3E, 3F**). Collectively, these results suggest that myofiber-specific ablation of IRE1α reduces the number of satellite cells in injured muscle microenvironment.

### Targeted ablation of IRE1 reduces proliferation of satellite cells in injured muscle

Using EdU labelling for proliferating cells, we sought to investigate whether myofiber-specific deletion of IRE1α affects satellite cell proliferation in a cell non-autonomous manner. TA muscle of *Ern1*^fl/fl^ and *Ern1*cKO mice were injured by intramuscular injection of 1.2% BaCl_2_ solution.

After 72 h, the mice were given a single intraperitoneal injection of EdU and the TA muscle was isolated 11 days later, and stained for the detection of EdU^+^ myonuclei. The boundaries of myofibers were identified by staining for laminin protein whereas nuclei were counterstained using DAPI (**Figure 4A**). Intriguingly, we found that that the number of EdU^+^ nuclei per myofiber was significantly reduced in TA muscle of *Ern1*cKO mice compared with corresponding *Ern1*^fl/fl^ mice (**Figure 4B**). Moreover, the proportion of myofibers containing two or more EdU^+^ centrally located nuclei was significantly reduced in *Ern1*cKO mice compared with *Ern1*^fl/fl^ mice (**Figure 4C**). Our analysis also showed that percentage of EdU^+^ nuclei in total nuclei was also significantly reduced in regenerating TA muscle of *Ern1*cKO mice compared with *Ern1*^fl/fl^ mice (**Figure 4D**). These results suggest that myofiber-specific deletion of IRE1α reduces proliferation and fusion of satellite cells with regenerating myofibers.

**FIGURE 4.**
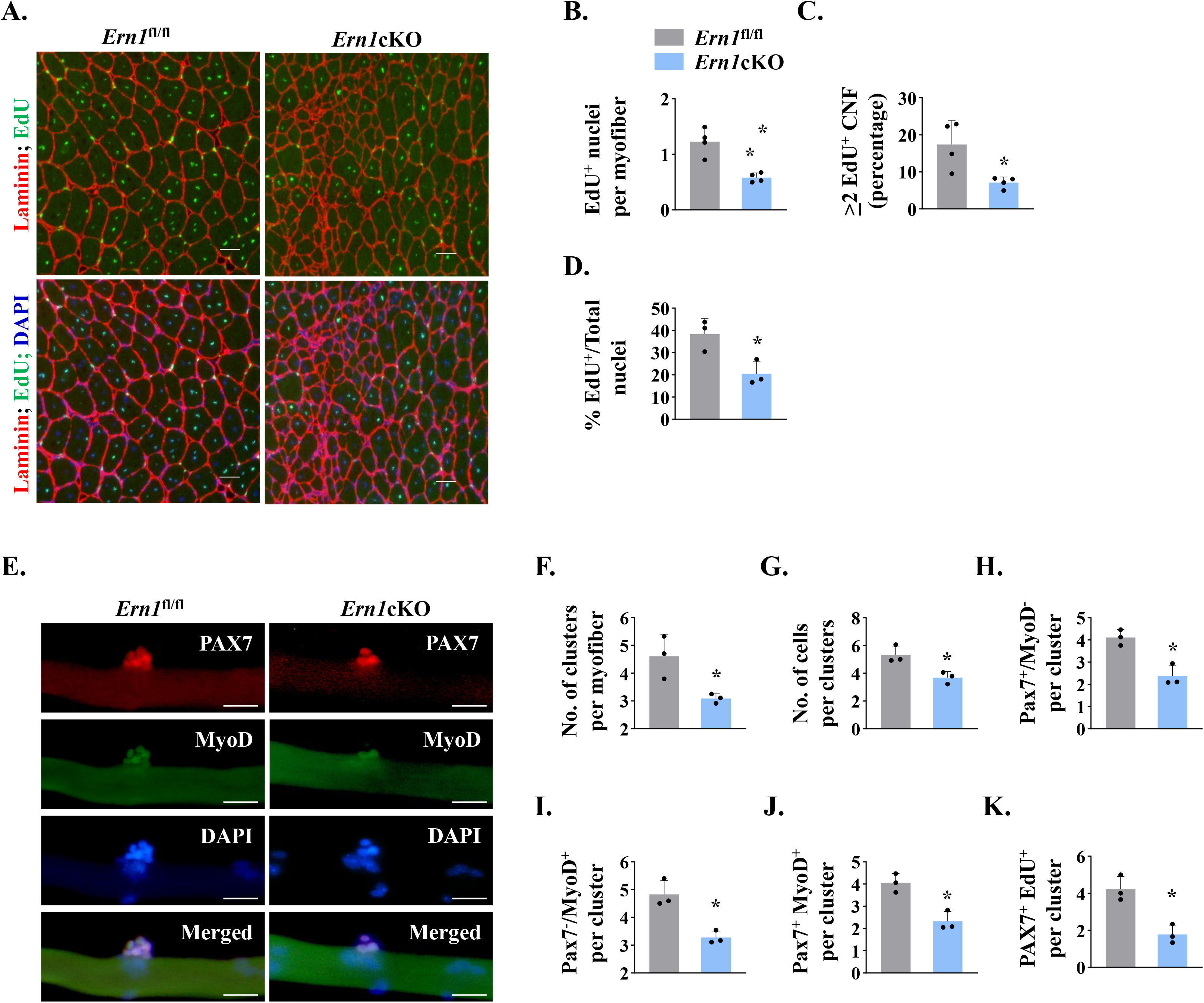
IRE1α promotes proliferation of satellite cells. (**A**) Left side TA muscle of *Ern1*^fl/fl^ and *Ern1*cKO mice was given intramuscular injection of 1.2% BaCl_2_ solution. After 3 days, the mice were given an intraperitoneal injection of EdU and 11 days later, the TA muscles were collected and muscle sections prepared were stained to detect EdU, laminin, and nuclei. Representative photomicrographs of TA muscle sections after EdU, anti-laminin, and DAPI staining are presented here. Scale bar: 50 µm. (**B**) Average number of EdU^+^ nuclei per myofiber, (**C**) percentage of myofibers containing 2 or more centrally located EdU^+^ nuclei (**D**) percentage of EdU^+^ nuclei to total nuclei in TA muscle of *Ern1*^fl/fl^ and *Ern1*cKO mice (n=4 mice per group). Data are presented as mean ± SEM. **\***P≤0.05, values significantly different from corresponding TA muscle of *Ern1*^fl/fl^ mice by unpaired t test. (**E**) Single myofibers were isolated from the EDL muscle of *Ern1*^fl/fl^ and *Ern1*cKO mice and cultured for 72 h. Representative images of myofiber-associated cells after immunostaining for Pax7 (red) and MyoD (green) protein. Nuclei were counterstained with DAPI (blue). Scale bar: 20 μm. Average number of (**F**) cellular clusters per myofiber, (**G**) cells per cluster, (**H**) Pax7^+/^MyoD^-^ cells per cluster, (**I**) Pax7^-/^MyoD^+^ cells per cluster, and (**J**) Pax7^+^/MyoD^+^ cells per cluster in *Ern1*^fl/fl^ and *Ern1*cKO cultures. (**K**) In a separate experiment, single myofibers after being cultured for 72 h were pulse labelled with EdU for 90 min and then immunostained for Pax7 protein and detection of EdU incorporation. Average number of Pax7^+^/EdU^+^ cells per cluster on *Ern1*^fl/fl^ and *Ern1*cKO myofibers (n=3 mice per group). Data are presented as mean ± SEM. **\***P≤0.05, values significantly different from myofiber cultures prepared from *Ern1*^fl/fl^ mice.

We also studied whether IRE1α influences the self-renewal or differentiation of satellite cells in a cell non-autonomous manner. We employed a suspension culture of EDL myofiber explants that represents an *ex vivo* model representing muscle injury *in vivo* with respect to the activation, proliferation, and differentiation of satellite cells (Hindi and Kumar, 2016; Xiong et al., 2017). In this system, each freshly isolated myofiber is associated with a fixed number quiescent (Pax7^+^/MyoD^−^) satellite cells. About 24h in cultures, satellite cells upregulate *Myod1* (Pax7^+^/MyoD^+^) and start proliferating to form cell aggregates on myofibers. Cells on cultured myofiber then either terminally differentiate (Pax7^-^/MyoD^+^) or undergo self-renewal (Pax7^+^/MyoD^-^) and enter quiescence (Hindi and Kumar, 2016; Ogura et al., 2015).

Immunostaining of freshly isolated myofibers from EDL muscle of *Ern1*^fl/fl^ and *Ern1*cKO mice showed no significant difference in the numbers of (Pax7^+^/MyoD^-^) cells and with undetectable levels of *Myod1* expression (data not shown). After 72h in suspension culture, the number of clusters per myofiber and number of cells per cluster were found to be significantly reduced in myofiber cultures prepared from *Ern1*cKO mice compared with *Ern1*^fl/fl^ mice (**Figure 4E-G**). Our analysis also showed that there was a significant decrease in number of self-renewing (Pax7^+^/MyoD^-^), proliferating (Pax7^+^/MyoD^+^), and differentiating (Pax7^-^/MyoD^+^) satellite cells per myofiber in *Ern1*^fl/fl^ cultures compared to *Ern1*cKO mice (**Figure 4H-J**) suggesting that deletion of IRE1α in myofiber reduces the overall pool of satellite cells without influencing their self-renewal, proliferation, or differentiation properties. To directly investigate the role of IRE1α in the proliferation of myofiber-associated satellite cells, we pulse-labelled satellite cells with EdU followed by immunostaining for Pax7 and detection of EdU (**Figure 4-figure supplement 1**). Results showed that the number of Pax7^+^/EdU^+^ satellite cells per cluster was significantly reduced in *Ern1*cKO mice compared with *Ern1*^fl/fl^ mice (**Figure 4K**).

### IRE1α improves skeletal muscle regeneration through XBP1

IRE1α is an endonuclease that causes the alternative splicing of *Xbp1* mRNA to generate a transcriptionally active XBP1 (sXBP1). In addition, IRE1α activation can cause the cleavage of other ER-localized mRNAs, cytosolic mRNAs, and microRNAs, leading to their degradation through a process named RIDD (Hollien and Weissman, 2006; Maurel et al., 2014). By performing qPCR, we first compared the levels of various mRNAs that are known to be degraded by RIDD process in uninjured and injured TA muscle of *Ern1*^fl/fl^ and *Ern1*cKO mice. Results showed that there was no significant difference in mRNA levels of known RIDD targets such as *Myod1*, *Hgsnat*, *ERdj4*, *Col6*, *Pdqfrb*, *Scara3*, and *Sparc* in uninjured or 5d-injured TA muscle of *Ern1*^fl/fl^ and *Ern1*cKO mice. A recent study showed that IRE1α reduces the levels myostatin in muscle cells through RIDD (He et al., 2021). However, we did not find any significant difference in mRNA levels of myostatin (*Mtsn*) between uninjured or 5d-injured TA muscle of *Ern1*^fl/fl^ and *Ern1*cKO mice (**Figure 5-figure supplement 1**). By contrast, mRNA levels of *sXbp1*were found to be drastically reduced in TA muscle of *Ern1*cKO mice compared with *Ern1*^fl/fl^ mice (**Figure 5-figure supplement 1A**).

To understand whether IRE1α promotes satellite cell proliferation and skeletal muscle regeneration in adult mice through activating XBP1 transcription factor, we crossed floxed *Xbp1* (*Xbp1*^fl/fl^) mice with *Mck*-Cre line to generate muscle-specific *Xbp1*-knockout (*Xbp1*cKO) and littermate *Xbp1*^fl/fl^ mice. There is a drastic reduction in the levels of both unspliced XBP1 and sXBP1 in skeletal muscle of *Xbp1*cKO mice compared with littermate *Xbp1*^fl/fl^ mice, as described (Bohnert et al., 2019; Parveen et al., 2021). Finally, TA muscle of littermate *Xbp1*^fl/fl^ and *Xbp1*cKO mice was injured through intramuscular injection of 1.2% BaCl_2_ solution and muscle regeneration was studied at day 5 and 14 days post-injury. H&E staining of TA muscle sections showed that there was a considerable reduction in the regeneration of TA muscle of *Xbp1*cKO mice compared to *Xbp1*^fl/fl^ mice (**Figure 5A**). Morphometric analysis also showed that there was a significant decrease in the average myofiber CSA and Minimal Feret’s diameter and proportion of myofibers containing two or more centrally located nuclei in 5d-injured TA muscle of *Xbp1*cKO mice compared with *Xbp1*^fl/fl^ mice (**Figure 5B-D**). Moreover, the size of the regenerating myofibers was also found to be significantly reduced in 14d-injured TA muscle of *Xbp1*cKO mice compared with *Xbp1*^fl/fl^ mice (**Figure 5A, 5E, 5F**).

**FIGURE 5.**
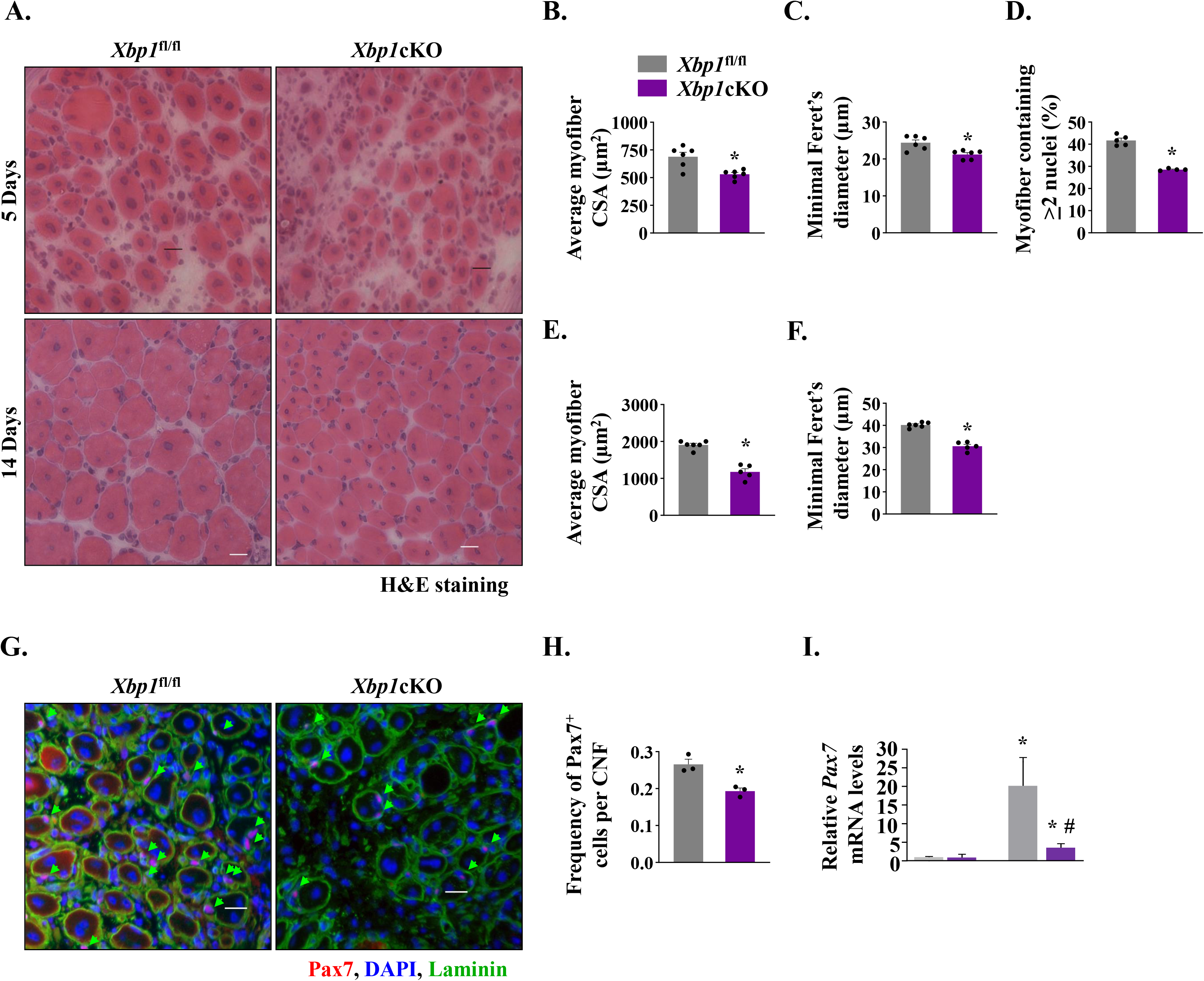
Myofiber-specific ablation of XBP1 in mice inhibits skeletal muscle regeneration. Left side TA muscle of *Xbp1*^fl/fl^ and *Xbp1*cKO mice was injured by intramuscular injection of 50 µl of 1.2% BaCl_2_ solution whereas right side TA muscle was injected with saline and served as control. The muscles were harvested at day 5 and 14 post-BaCl_2_ injection. (**A**) Representative photomicrographs of H&E-stained sections of 5d- and 14d-injured TA muscle of *Xbp1*^fl/fl^ and *Xbp1*cKO mice. Scale bar: 20 µm. Quantification showing (**B**) Average myofiber CSA (n=6 mice per group) (**C**) average Minimal Feret’s diameter (n=6 mice per group), and (**D**) Percentage of myofibers containing 2 or more centrally located nuclei in TA muscle of *Xbp1*^fl/fl^ and *Xbp1*cKO mice at day 5 post-injury. Quantification of average (**E**) myofiber CSA and (**F**) minimal Feret’s diameter in TA muscle of *Xbp1*^fl/fl^ and *Xbp1*cKO mice at day 14 post-injury. (n=5 for Injured *Xbp1*^fl/fl^ and n=4 for Injured *Xbp1*cKO group) (**G**) Representative photomicrographs of 5d-injured TA muscle sections from *Xbp1*^fl/fl^ and *Xbp1*cKO mice after immunostaining for Pax7 (red) and laminin (green) protein. Nuclei were stained with DAPI (blue). Scale bar: 20 µm. (**H**) Average number of Pax7^+^ cells per centrally nucleated myofiber in 5d-injured TA muscle of *Xbp1*^fl/fl^ and *Xbp1*cKO mice n=4 mice per group. Data are presented as mean ± SEM. **\***P≤0.05, values significantly different from *Xbp1*^fl/fl^ mice. (**I**) Relative levels of *Pax7* mRNA in uninjured and 5d-injured TA muscle of *Xbp1*^fl/fl^ and *Xbp1*cKO mice. Data are presented as mean ± SEM and analyzed by one-way analysis of variance (ANOVA) followed by Tukey’s multiple comparison test. **\***P≤0.05, values significantly different from uninjured TA muscle of *Xbp1*^fl/fl^ mice. ^#^P ≤ 0.05, values significantly different from 5d-injured TA muscle of *Xbp1*^fl/fl^ mice.

By performing immunostaining for Pax7 protein, we also quantified the number of satellite cells in *Xbp1*^fl/fl^ and *Xbp1*cKO mice. There was no significant difference in the number of Pax7^+^ cells in uninjured TA muscle of *Xbp1*^fl/fl^ and *Xbp1*cKO mice. However, the abundance of satellite cells was found to be significantly reduced in 5d-injured TA muscle of *Xbp1*cKO mice compared with *Xbp1*^fl/fl^ mice (**Figure 5G, H**). Moreover, mRNA levels of *Pax7* were also found to be significantly reduced in 5d-injured TA muscle of *Xbp1*cKO mice compared with *Xbp1*^fl/fl^ mice (**Figure 5I**). While we observed a significant reduction in the number of Pax7^+^ cells in regenerating TA muscle of *Xbp1*cKO mice, it remains unknown whether this is due to reduced proliferation or survival of satellite cells. Future studies will determine how XBP1 signaling within myofibers affect the satellite cell number in injured muscle microenvironment. Altogether, these results suggest that IRE1α promotes skeletal muscle regeneration through the activation of XBP1 transcription factor.

### IRE1α regulates Notch signaling in injured muscle microenvironment

To understand the mechanisms by which IRE1α-mediated signaling promotes proliferation of satellite cells, we first studied the gene expression of insulin growth factor-1 (*Igf1*), fibroblast growth factor (*Fgf*) 1 and 2, hepatocyte growth factor (*Hgf*), and stromal-derived factor 1 (*Sdf1*) which are known to promote satellite cell proliferation (Yin et al., 2013). We found no significant difference in mRNA levels of *Igf1*, *Fgf1*, *Hgf*, or *Sdf1* in 5d-injured TA muscle of *Ern1*^fl/fl^ and *Ern1*cKO mice. There was a small but significant increase in mRNA levels of *Fgf2* in 5d-injured muscle of *Ern1*cKO mice compared with littermate *Ern1*^fl/fl^ mice (**Figure 6-figure supplement 1**).

Several studies have demonstrated that Notch signaling is essential for the self-renewal and proliferation of satellite cells during regenerative myogenesis. To further understand the potential mechanisms through which IRE1α signaling in myofibers regulate satellite cell proliferation and skeletal muscle regeneration, we first measured mRNA levels of Notch ligands (*Jagged1*, *Jagged2*, *Dll1* and *Dll4*), Notch receptors (*Notch1, Notch2*, and *Notch3*) and target genes (*Hes1, Hes6, Hey1* and *Heyl*) by qPCR assay. Interestingly, mRNA levels of *Notch1*, *Notch2*, *Notch3*, *Jagged1*, *Jagged2*, *Hes6*, *Hey1*, and *Heyl* were found to be significantly reduced in 5d-injured TA muscle of *Ern1*cKO mice compared with *Ern1*^fl/fl^ mice (**Figure 6A-C)**. Western blot analysis also showed that the protein levels of Notch1 and Hes6 were considerably reduced in injured TA muscle of *Ern1*cKO mice compared with *Ern1*^fl/fl^ mice (**Figure 6D**). These results suggest that myofiber-specific ablation of IRE1α reduces the activation of Notch signaling during skeletal muscle regeneration.

**FIGURE 6.**
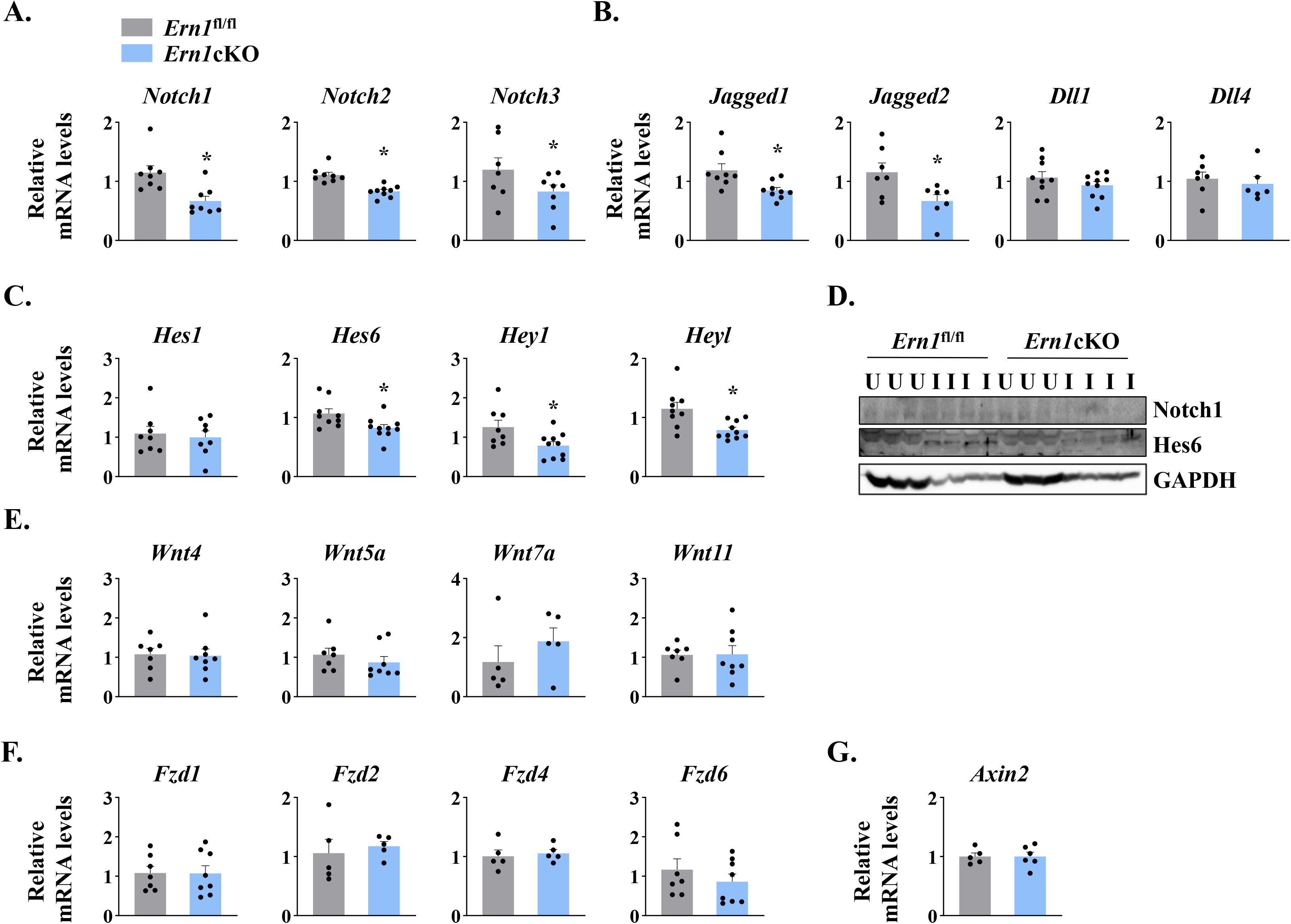
IRE1α regulates Notch signaling during skeletal muscle regeneration. TA muscle of *Ern1*^fl/fl^ and *Ern1*cKO mice were injected with 1.2% BaCl_2_ solution. After 5d, the muscles were isolated and processed for qPCR and western blot analysis. Relative mRNA levels of (**A**) Notch receptors *Notch1*, *Notch2*, and *Notch3*; (**B**) Notch ligands *Jagged1*, *Jagged2*, *Dll1*, and *Dll4*; and (**C**) Notch targets *Hes1*, *Hes6*, *Hey1* and *Heyl* in 5d-injured TA muscle of *Ern1*^fl/fl^ and *Ern1*cKO mice (n=8-9 per group). (**D**) Immunoblot presented here demonstrate protein levels of Notch1, Hes6, and unrelated protein GAPDH in uninjured and 5d-injured TA muscle of *Ern1*^fl/fl^ and *Ern1*cKO mice. Relative mRNA levels of (**E**) Wnt ligands *Wnt4*, *Wnt5A*, *Wnt7A* and *Wnt11*; (**F**) Wnt receptors *Fzd1*, *Fzd2*, *Fzd4*, and *Fzd6*, and (**G**) Wnt targets *Axin2* in 5d-injured TA muscle of *Ern1*^fl/fl^ and *Ern1*cKO mice (n=5 per group). Data are presented as mean ± SEM. **\***P≤0.05, values significantly different from corresponding 5d-injured TA muscle of *Ern1*^fl/fl^ mice.

Wnt signaling also plays an important role during skeletal muscle regeneration. The levels of various Wnt ligands and receptors are increased in skeletal muscle upon injury. Moreover, Wnt signaling has been found to promote myoblast fusion both *in vivo* and *in vitro*. We investigated whether targeted ablation of IRE1α influences the activation of Wnt pathway during skeletal muscle regeneration. However, we found no significant difference in the gene expression of Wnt ligands *Wnt4*, *Wnt5a*, *Wnt7a* and *Wnt11*, Wnt receptors *Fzd1*, *Fzd2*, *Fzd4* and *Fzd6*, and Wnt target gene *Axin2* (**Figure 6E-G**). These results suggest that IRE1α specifically regulate Notch signaling to promote satellite cell proliferation.

### IRE1α regulates NF-κB signaling in regenerating skeletal muscle

NF-κB is a major transcription factor that regulates the expression of a plethora of molecules involved in cell proliferation, survival, and differentiation and inflammatory response (Hayden and Ghosh, 2004; Razani et al., 2011). NF-κB signaling has also been found to play an important role in skeletal muscle regeneration. There are multiple reports suggesting a crosstalk between Notch and NF-κB signaling in diverse experimental model. Indeed, Notch1 has been shown to increase the expression of various subunits of NF-κB complex. Notch1 intracellular domain (N1ICD) activates canonical NF-κB pathway through physically interacting with IKK signalosome and through repressing the deubiquitinase CYLD, a negative IKK complex regulator (Espinosa et al., 2010; Ferrandino et al., 2018; Osipo et al., 2008). By performing western blot, we first measured levels of phosphorylated and total p65 protein (a marker for activation of canonical NF-κB signaling). A drastic increase in the levels of both phosphorylated p65 (p-p65) and total p65 was observed in 5d-injured TA muscle of both *Ern1*^fl/fl^ and *Ern1*cKO mice. However, the levels of p- p65 protein were found to be significantly reduced in 5d-injured TA muscle of *Ern1*cKO mice compared with *Ern1*^fl/fl^ mice (**Figure 7A, 7B**). We also measured the levels of p100 and p52, the markers of activation of non-canonical NF-κB signaling pathway. Muscle injury drastically increased the levels of p100 and p52 protein in TA muscle of both *Ern1*^fl/fl^ and *Ern1*cKO mice. Intriguingly, protein levels of both p100 and p52 were found to be significantly higher in injured TA muscle of *Ern1*cKO mice compared with *Ern1*^fl/fl^ mice (**Figure 7A, 7C**).

**FIGURE 7.**
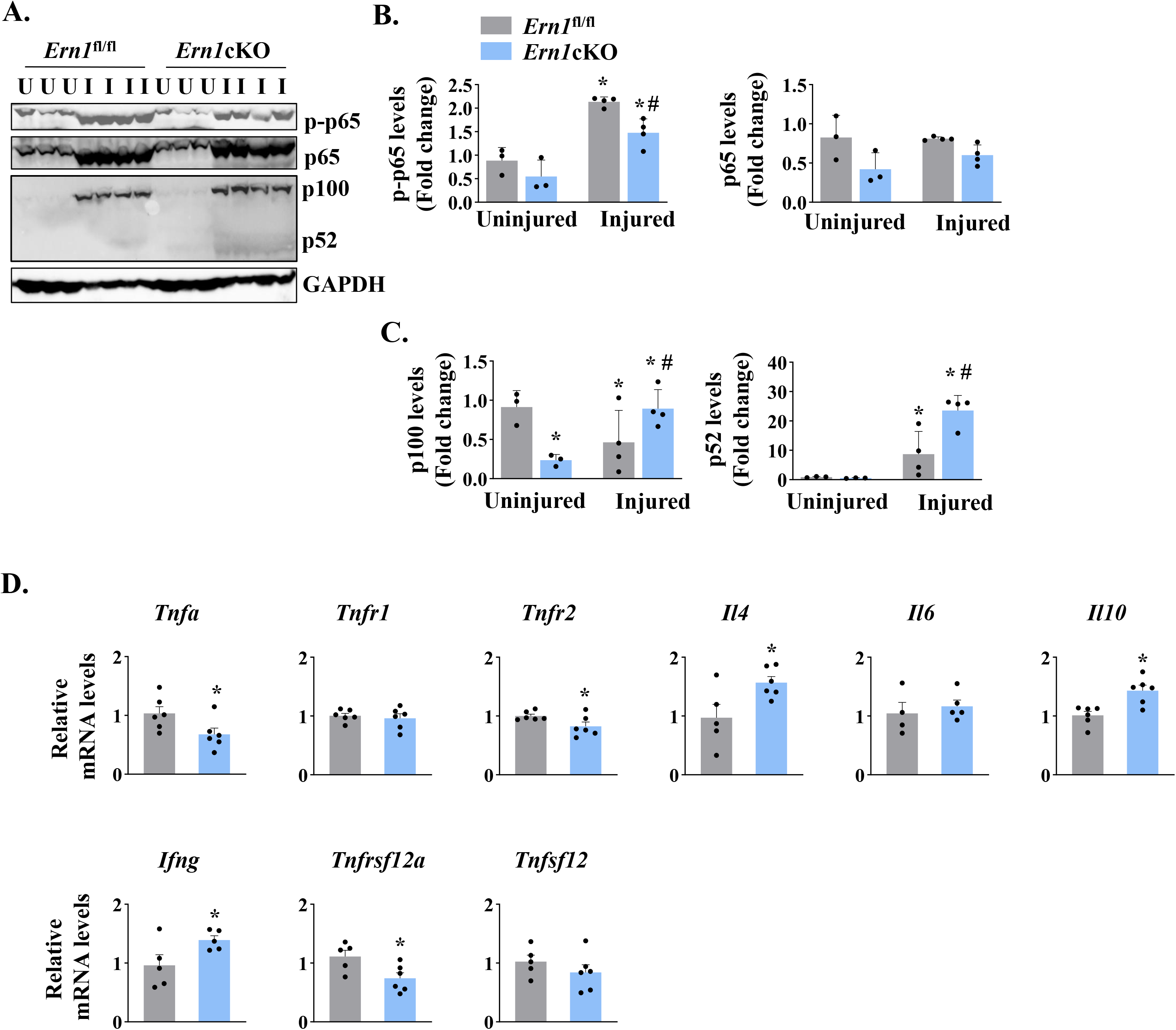
IRE1α regulates NF-κB signaling in regenerating myofibers. Left side TA muscle of *Ern1*^fl/fl^ and *Ern1*cKO mice was given intramuscular injection of 1.2% BaCl_2_ solution whereas right side TA muscle was injected with saline alone. After 5 days, the TA muscles were harvested and processed by western blotting and qPCR. (**A**) Representative immunoblots presented her demonstrate the levels of p-p65, p65, p100, p52 and unrelated protein GAPDH in uninjured and 5d-injured TA muscle of *Ern1*^fl/fl^ and *Ern1*cKO mice. Quantification of levels of (**B**) p-p65 and p65 (**C**) p100 and p52 protein (n=3 or 4 mice per group). Data are presented as mean ± SEM and analyzed by one-way analysis of variance (ANOVA) followed by Tukey’s multiple comparison test. **\***P≤0.05, values significantly different from uninjured TA muscle of *Ern1*^fl/fl^ mice. ^#^P ≤ 0.05, values significantly different from 5d-injured TA muscle of *Ern1*^fl/fl^ mice. (**D**) Relative mRNA levels of *Tnfa*, *Tnfr1*, *Tnfr2*, *Il4*, *Il6*, *Il10, Ifng, Tnfsf12*, *Tnfrsf12a* in 5d-injured TA muscle of *Ern1*^fl/fl^ and *Ern1*cKO mice (n=6 mice per group). Data are presented as mean ± SEM. **\***P≤0.05, values significantly different from 5d-injured TA muscle of *Ern1*^fl/fl^ mice. U, uninjured; I, injured.

We also measured mRNA levels of a few cytokines that regulate myogenesis. Our qPCR analysis showed that mRNA levels of *Tnfa*, *Tnfr2*, and *Tnfrsf12a* were found to be significantly reduced in injured TA muscle of *Ern1*cKO mice compared with *Ern1*^fl/fl^ mice (**Figure 7D**). By contrast, mRNA levels of *Il4*, *Il10, and Ifng* were significantly increased in injured TA muscle of *Ern1*cKO mice compared to *Ern1*^fl/fl^ mice. There was no significant difference in the mRNA levels of *Tnfr1*, *Tnfsf12*, and *Il6* between injured muscle of *Ern1*^fl/fl^ and *Ern1*cKO mice (**Figure 7D**). Taken together, these results suggest that myofiber-specific ablation of IRE1α influences the activation of canonical and non-canonical NF-κB pathways and regulates the expression of key cytokines and their receptors during regenerative myogenesis.

### Myofiber-specific deletion of IRE1 exacerbates dystrophic phenotype in mdx mice

In the preceding experiments, we used a mouse model that involves single acute injury to TA muscle followed by its regeneration. However, the role of IRE1 signaling during chronic muscle injury is not known. Dystrophin-deficient myofibers undergo chronic muscle injury and regeneration. The mdx mouse, which lacks dystrophin protein due to a mutation that results in a premature stop codon in exon 23, is widely used as a mouse model for studying chronic muscle injury and pathophysiology of Duchenne muscular dystrophy (Chang et al., 2016; Shin et al., 2013). Thus, we employed mdx mice to evaluate the role of IRE1α in muscle regeneration in the “settings” of chronic muscle injury and myopathy.

We first investigated how IRE1/XBP1α pathway is affected in skeletal muscle of mdx mice. Results showed that the levels of phosphorylated IRE1α (p-IRE1α) was significantly increased whereas total IRE1α protein was significantly reduced in skeletal muscle of mdx mice compared to corresponding control mice. Indeed, the ratio of phosphorylated versus total IRE1α as well as levels of sXBP1 protein were significantly higher in skeletal muscle of mdx mice compared with control mice (**Figure 8A, 8B**) suggesting activation of IRE1α/XBP1 pathway in dystrophic muscle of mdx mice. To understand the role of IRE1α in regeneration of dystrophic muscle, we crossed *Ern1*cKO mice with mdx mice to generate littermate mdx;*Ern1*^fl/fl^ and mdx;*Ern1*cKO mice. There was no significant difference in the body weight of 12-week old littermate mdx;*Ern1*^fl/fl^ and mdx;*Ern1*cKO mice (**Figure 8C**). However, there was a significant reduction in four paw grip strength (normalized with body weight) of mdx;*Ern1*cKO mice compared with mdx;*Ern1*^fl/fl^ mice (**Figure 8D**). We next isolated hind limb muscle from these mice and performed H&E staining (**Figure 8E**). Interestingly, our results showed that average CSA of the regenerating myofiber (centronucleated) was significantly reduced in TA muscle of mdx;*Ern1*cKO mice compared to mdx;*Ern1*^fl/fl^ mice (**Figure 8E, 8F**). Furthermore, there was a significant reduction in number of myofibers containing two or more centrally located nuclei in mdx;*Ern1*cKO mice compared with mdx;*Ern1*^fl/fl^ mice (**Figure 8G**). Similarly, muscle regeneration was also considerably reduced in GA muscle of *Ern1*cKO mice compared with mdx;*Ern1*^fl/fl^ mice (**Figure 8-figure supplement 1A, 1B**). We also investigated how myofiber-specific deletion of IRE1α affects the frequency of satellite cells in dystrophic muscle of mdx mice. TA muscle sections generated from littermate mdx;*Ern1*^fl/fl^ and mdx;*Ern1*cKO mice were immunostained for Pax7 and laminin whereas nuclei were counterstained with DAPI. Finally, the number of Pax7^+^ cells within laminin staining was counted. Results showed that frequency of satellite cells per myofiber was significantly reduced in TA muscle of mdx;*Ern1*cKO mice compared with mdx;*Ern1*^fl/fl^ mice (**Figure 8H, 8I**). There was also a significant reduction in the mRNA levels of *Pax7* in TA muscle of mdx;*Ern1*cKO mice compared with mdx;*Ern1*^fl/fl^ mice (**Figure 8J**). Finally, our qPCR analysis showed that there was no significant difference in mRNA levels *Bloc1s1* or *Mtsn* (myostatin) in dystrophic muscle of mdx;*Ern1*^fl/fl^ and mdx;*Ern1*cKO mice (**Figure 8-figure supplement 1C**). By contrast, mRNA levels of Notch targets *Hes1*, *Hey1*, and *Heyl* were found to be significantly reduced in skeletal muscle of mdx;*Ern1*cKO mice compared with mdx;*Ern1*^fl/fl^ mice (**Figure 8-figure supplement 1D**). These results further suggest that IRE1α improves muscle regeneration through augmenting satellite cell proliferation in response to acute injury and in dystrophic muscle of mice.

**FIGURE 8.**
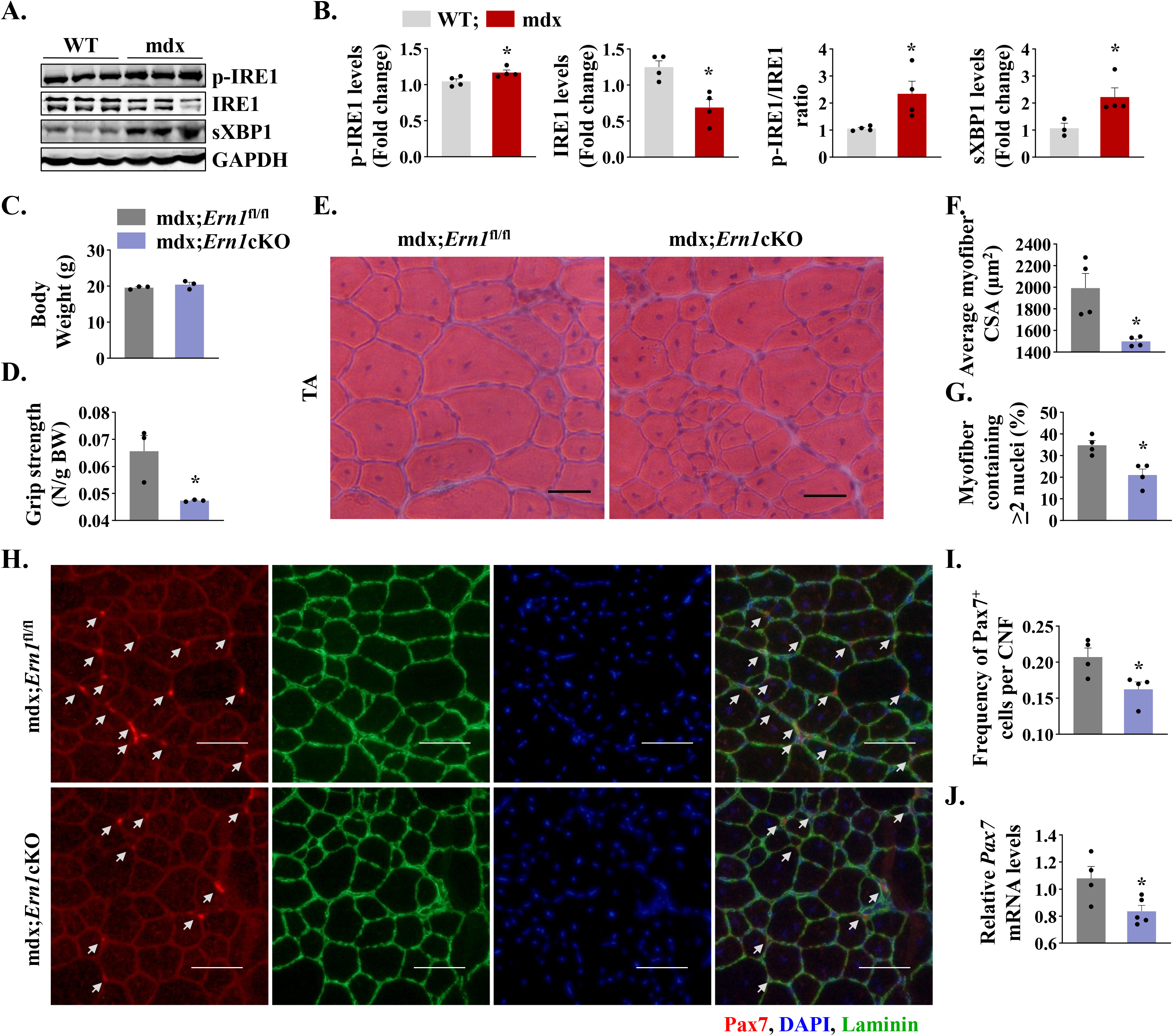
Myofiber-specific ablation of IRE1α exacerbates myopathy in mdx mice. (**A**) Western blots showing the levels of p-IRE1, IRE1, sXBP1 and unrelated protein GAPDH in TA muscle of 12-week old wild-type (WT) and mdx mice. (**B**) Densitometry analysis showing levels of p-IRE1, IRE1, sXBP1 and GAPDH protein in TA muscle of wild-type and mdx mice (n=3 in each group). *P<0.05, values significantly different from WT mice. (**C**) Average body weight (BW), and (**D**) Four limb grip strength normalized with body weight of 10-week old mdx;*Ern1*^fl/fl^ and mdx;*Ern1*cKO mice (n=3 in each group). (**E**) Representative photomicrographs of H&E-stained TA muscle section from 10-week old mdx;*Ern1*^fl/fl^ and mdx;*Ern1*cKO mice. Scale bar: 50 µm. (**F**) Average myofiber CSA and (**G**) percentage of myofibers containing ≥2 nuclei in TA muscle of mdx;*Ern1*^fl/fl^ and mdx;*Ern1*cKO mice (n=4 in each group). (**H**) Representative photomicrographs of TA muscle sections after immunostaining for Pax7 (red) and laminin (green) protein. Nuclei were stained with DAPI (blue). Scale bar: 50 µm. White arrows point to Pax7^+^ satellite cells. (**I**) Frequency of Pax7^+^ cells per centrally nucleated myofibers in TA muscle of mdx;*Ern1*^fl/fl^ and mdx;*Ern1*cKO mice (n=4 in each group). (**J**) Relative mRNA levels of *Pax7* in TA muscle of mdx;*Ern1*^fl/fl^ and mdx;*Ern1*cKO mice assayed by performing qPCR (n=5 in each group). Data are presented as mean ± SEM. **\***P≤0.05, values significantly different from TA muscle of mdx;*Ern1*^fl/fl^ mice.

## Discussion

Skeletal muscle repair in response to injury is regulated by multiple factors produced by damaged myofibers, satellite stem cells, and several other cell types that are either resident in the muscle or recruited to assist in clearing cellular debris and support regeneration (Yin et al., 2013). Furthermore, it is now increasingly clear that several signaling pathways are activated in injured myofibers that regulate specific steps of muscle regenerative program in adults (Brack and Munoz-Canoves, 2016; Dumont et al., 2015). Recent studies have provided evidence that many catabolic stimuli including muscle injury lead to the activation of ER stress-induced UPR pathways in skeletal muscle (Afroze and Kumar, 2019; Bohnert et al., 2018; Miyake et al., 2016; Xiong et al., 2017). In the present study, we demonstrate that muscle injury leads to the activation of IRE1/XBP1 signaling axis in skeletal muscle and that deletion of IRE1 or XBP1 in myofibers attenuate skeletal muscle regeneration potentially through inhibiting the proliferation of satellite cells in cell non-autonomous manner. Our study also demonstrates that IRE1- mediated signaling in myofiber is essential for the regeneration of injured myofibers in dystrophic muscles of mdx mice.

Skeletal muscle regeneration is accomplished through the activation of satellite cells which proliferate and differentiate to become myoblasts that eventually fuse with injured myofibers to complete regeneration (Kuang and Rudnicki, 2008; Yin et al., 2013). While the frequency of satellite cells in skeletal muscle of healthy individuals remain constant, their number and regenerative potential can be altered in various disease states and conditions (Brack and Munoz-Canoves, 2016; Dumont et al., 2015). We found that ablation of IRE1α in myofibers has no noticeable effect on skeletal muscle development or myofiber size in mice. Furthermore, genetic ablation of IRE1α in differentiated myofibers does not influence the frequency of satellite cells in naïve conditions (**Figure 3**). In contrast, myofiber-specific deletion of IRE1α inhibits the regeneration of skeletal muscle in response to injury in adult mice. Our results also demonstrate that the size of the regenerating myofibers and levels of various MRFs, such as *Myf5*, *Myod1* and *Myog* as well as *Myh3* are considerably reduced in injured muscle of *Ern1*cKO mice (**Figure 2**) suggesting that IRE1α-mediated signaling supports the early steps of muscle regenerative program. We have previously reported that PERK is essential for the survival and differentiation of activated satellite cells into the myogenic lineage. While the role of IRE1 in muscle progenitor cells has not been yet investigated, targeted ablation of XBP1 does not affect satellite cell function during regenerative myogenesis (Xiong et al., 2017). These results suggest that distinct arms of the UPR may have different roles in the regulation of muscle progenitor cell function and skeletal muscle regeneration in adults. It is also possible that IRE1 regulates satellite cell function through its kinase activity which will be investigated in future studies.

IRE1α, the most evolutionary conserved UPR signaling branch, is a serine/threonine protein kinase and endoribonuclease that cleave and initiate splicing of the *Xbp1* mRNA (Tabas and Ron, 2011; Walter and Ron, 2011). Activated IRE1α endoribonuclease removes 26-nucleotide intron from *Xbp1*, resulting in a translational frame-shift to modify unspliced XBP1 into spliced XBP1 transcription factor that enforces adaptive programs (Hetz, 2012; Walter and Ron, 2011; Wang and Kaufman, 2014). Activated IRE1α also controls various biological processes, including cell death and inflammation through degradation of a subset of mRNAs and microRNAs through a process known as RIDD (Hollien and Weissman, 2006; Maurel et al., 2014). The target mRNAs of RIDD in mammalian cells require a consensus motif of 5′- CUGCAG-3′, along with a secondary stem loop structure. The disruption in stem loop formation or mutation in the consensus sequence hampers RIDD-mediated degradation of the target mRNA (Bright et al., 2015; Oikawa et al., 2010). Intriguingly, in response to some stimuli, such as genotoxic stress or microbial products, IRE1α signaling is activated without any noticeable ER stress signature (Abdullah and Ravanan, 2018; Dufey et al., 2020; Martinon et al., 2010). For example, in response to DNA damage in fibroblasts, IRE1α signaling is activated in the absence of an ER stress leading to the activation of RIDD process without having any effect on *Xbp1* mRNA splicing (Dufey et al., 2020). There is also a recent report suggesting that IRE1α promotes skeletal muscle regeneration through reducing the levels of myostatin via RIDD mechanism (He et al., 2021). Although we have used different floxed *Ern1* mice and employed *Mck*-Cre line to ablate IRE1α in myofibers, we did not observe any significant difference in the levels of myostatin in uninjured or injured muscle of *Ern1*^fl/fl^ and *Ern1*cKO mice (**Figure 5- figure supplement 1**). Furthermore, there was no difference in the levels of other mRNAs which are known to be degraded through RIDD-dependent mechanisms suggesting that IRE1α signaling in myofibers may not be regulating skeletal muscle regeneration through RIDD pathway (**Figure 5-figure supplement 1)**. Our results are consistent with the findings that the RIDD-dependent mRNA decay occurs only when there is an ectopic activation of IRE1α with concomitant inhibition of protein synthesis (Moore and Hollien, 2015). Indeed, our results demonstrating that myofiber-specific ablation of *Xbp1* using the same *Mck*-Cre line produces similar effects as that of IRE1α (**Figure 5**) suggest that IRE1α promotes skeletal muscle regeneration through activating its canonical output mediated by the transcription factor XBP1. However, we cannot rule out the possibility that muscle progenitors also require RIDD or kinase activity of IRE1 for regeneration of injured skeletal muscle as has been observed in the recently published report where IRE1 was deleted specifically in myoblasts using *Myod1*-Cre line (He et al., 2021). Certainly, more investigations are needed to understand the mechanisms of action of IRE1α during regenerative myogenesis.

Strikingly, our results demonstrate that genetic ablation of IRE1 or XBP1 in myofibers significantly reduces the proliferation of satellite cells without having any significant impact on their self-renewal, survival, or differentiation (**Figures 3, 4, 5**). Notch signaling is quite unique that it can regulate the fate of one cell with that of a cellular neighbor through interaction between the Notch receptor and the membrane-bound Notch ligands that are expressed in a juxtaposed cell (Kann et al., 2021). It is now evidenced that the outcome of Notch signaling is dependent on the cellular context and is capable of influencing quiescence, self-renewal, survival, and differentiation cell fates. Notch signaling has also been shown to regulate the self-renewal and proliferation of satellite stem cells both *in vivo* and *in vitro* (Bi et al., 2016; Kann et al., 2021; Vasyutina et al., 2007). Interestingly, our results demonstrate that levels of Notch receptor 1, 2 and 3 and Notch ligands Jagged1 and Jagged2 are significantly reduced in injured muscle of *Ern1*cKO mice compared with *Ern1*^fl/fl^ mice. In addition, transcript levels of Notch target genes are also reduced in regenerating muscle of *Ern1*cKO mice further suggesting an inhibition in Notch signaling upon myofiber-specific ablation of IRE1α (**Figure 6**). While we observed an inhibition in the markers of Notch signaling in *Ern1*cKO mice, it remains unknown how depletion in IRE1 in myofibers influences Notch signaling. It is possible that IRE1/XBP1 signaling directly regulates the gene expression of Notch ligands in myofibers which through interaction with Notch receptor promotes the proliferation of other cell types in regenerating myofibers. However, it is also possible that IRE1-mediated signaling regulates Notch signaling through paracrine mechanisms. Certainly, more investigation is needed to understand the mechanisms by which IRE1/XBP1 arm of the UPR affects Notch signaling during regenerative myogenesis.

Previous studies have also shown that the levels of various components of canonical Wnt signaling are increased in injured skeletal muscle and Wnt signaling promotes myoblast fusion during regenerative myogenesis (Hindi et al., 2017; Hindi et al., 2013). Indeed, spatial activation of Notch and Wnt signaling is important for progression of muscle regeneration upon injury (Brack et al., 2008). However, there was no significant difference in the expression of various components of Wnt signaling in regenerating muscle of *Ern1*^fl/fl^ and *Ern1*cKO mice (**Figure 6**) suggesting that IRE1α specifically regulates Notch signaling and inhibition of Notch signaling may be one of the important mechanisms for the reduced proliferation of satellite cells in *Ern1*cKO mice.

There is a plethora of literature suggesting a crosstalk between Notch and NF-κB signaling pathways especially in various types of cancers. In general, Notch-1 and Notch target genes augments the activity of NF-κB through multiple mechanisms (Espinosa et al., 2010; Ferrandino et al., 2018; Osipo et al., 2008). NF-κB is known to promote proliferation and survival of a number of cell types. NF-κB can be activated through a canonical pathway that involves the activation of IKKβ and p65 subunits. In non-canonical pathway, protein levels of NIK are increased due to inhibition of its degradation. NIK phosphorylates IKKα which in turn phosphorylates p100 protein leading to its proteolytic processing into p52 subunit (Hayden and Ghosh, 2004; Razani et al., 2011). Accumulating evidence suggests that canonical NF-κB signaling is activated at initial stages of myogenesis to augment the proliferation of muscle progenitor cells (Li et al., 2008; Straughn et al., 2019). In contrast, non-canonical NF-κB signaling augments myoblast fusion during myogenesis (Enwere et al., 2012; Hindi et al., 2017; Hindi et al., 2013). Interestingly, our results suggest that the levels of phosphorylated p65 protein as well as gene expression of *Tnf*α, a potent inducer of canonical NF-κB signaling (Hayden and Ghosh, 2004), are significantly reduced in regenerating myofibers of *Ern1*cKO mice compared to *Ern1*^fl/fl^ mice. In contrast, there was a significant increase in the levels of p100 and p52 proteins in the injured muscle of *Ern1*cKO mice compared to *Ern1*^fl/fl^ mice suggesting activation of non-canonical NF-κB signaling upon deletion of IRE1α (**Figure 7**). Although the physiological significance of differential activation of canonical and non-canonical NF-κB pathway remains unknown, it is possible that inhibition of canonical signaling is another mechanism for the reduced proliferation of satellite cells in injured skeletal muscle of *Ern1*cKO mice. Alternatively, spurious activation of non-canonical NF-κB signaling may cause precautious fusion of muscle progenitor cells leading to deficits in muscle regeneration.

DMD is a genetic muscle disorder which involves chronic degeneration and regeneration of myofibers (Emery, 1998). It has been postulated that inadequate muscle regeneration in DMD is due to exhaustion of satellite cells after several rounds of myofiber degeneration and regeneration (Chang et al., 2016). In contrast to DMD patients, skeletal muscles of mdx mice maintain their ability to regenerate throughout life. Indeed, mdx mice serve as an excellent model to study muscle regeneration in disease conditions and in response to chronic myofiber injury (Blake et al., 2002; Chang et al., 2016; Shin et al., 2013). Our results demonstrate that genetic ablation of IRE1α exacerbates dystrophic phenotype and reduces the proportion of regenerating myofibers in skeletal muscle mdx mice (**Figure 8**). These results are consistent with the recently published report demonstrating that myoblast-specific deletion of IRE1α also exacerbates dystrophic phenotype in mdx mice (He et al., 2021). In contrast to this published report (He et al., 2021), we did not find any significant difference in the levels of myostatin or Bloc1S1 suggesting that IRE1α does not influence RIDD process in dystrophic muscle of mdx mice.

Interestingly, our analysis also showed that myofiber-specific deletion of IRE1α significantly reduced the number of satellite cells (**Figure 8**) and gene expression of Notch targets in skeletal muscle of mdx mice (**Figure 8-figure supplement 1D**) further suggesting that IRE1α signaling in myofibers promotes proliferation of satellite cells to support muscle regeneration. In summary, our study provides initial evidence that IRE1/XBP1 signaling in myofibers is activated during acute or chronic muscle injury and its main function is to promote muscle regeneration through augmenting proliferation of satellite cells in a cell non-autonomous manner. One caveat of our study is that we have used Cre-loxP system to delete *Ern1*or *Xbp1* gene in skeletal muscle of mice. While tissue-specific knockout is an important approach to understanding the function of a gene, it does not cause the complete excision of the target gene. Moreover, deletion of a gene from the beginning of the development can also lead to the activation of another gene to compensate for its functions. Therefore, more investigations using inducible *Ern1*- or *Xbp1*-knockout mice and pharmacological inhibitors are needed to establish the role of IRE1α/XBP1 axis in myofiber regeneration. Moreover, it will be important to determine whether supraphysiological activation of the IRE1α/XBP1 axis using transgenic or pharmacological approaches can improve myofiber regeneration in response to acute injury and in animal models of muscle degenerative diseases. Future studies should also investigate the cell-autonomous role of IRE1α in satellite cell function during regenerative myogenesis.

## Materials and Methods

| Key Resources Table |  |  |  |  |
| --- | --- | --- | --- | --- |
| Reagent type (species/strain or resource) | Designation | Source or reference | Research Resource Identifier (RRID) | Additional information |
| Genetic reagent (*Mus musculus*) | Wild type mice/C57BL/6 | The Jackson Laboratory | RRID:IMSR_JAX:000664 | Also known as B6, B6J, B6/J mice |
| Genetic reagent (*Mus musculus*) | Mdx mice/C57BL/10 ScSn<sup>-Dmdmdx</sup>/J | The Jackson Laboratory | RRID:IMSR_JAX:001801 | Also known as mdx mice |
| Genetic reagent (*Mus musculus*) | MCK-Cre Mice/Tg(Ckmm-cre)5Khn | The Jackson Laboratory | RRID:IMSR_JAX:006475 | Also known Ckmm-cre mice |
| Genetic reagent<br>(*Mus musculus*) | *Ern1*<sup>fl/fl</sup> Mice<br>/B6;129S4-<br>*Ern1*<sup>tm2.1Tiw</sup> | Kanazawa<br>Medical<br>University, Japan | RRID:IMSR_RBR<br>C05515 | Also known<br>as floxed<br>Ern1 mice |
| Genetic reagent<br>(*Mus musculus*) | *Xbp1*<sup>fl/fl</sup><br>mice/*Xbp1*<sup>tm2Glm</sup> | Dana-Farber<br>Cancer Institute,<br>Boston, MA | RRID:MGI:377401<br>7 | Also known<br>as floxed<br>Xbp1 mice |
| Antibody | anti-phospho-IRE1α<br>(rabbit<br>monoclonal) | Novus<br>Biologicals,<br>Cat# NB 100-2323 | RRID:AB_101452<br>03 | Dilution used<br>(1:500) |
| Antibody | anti-IRE1α<br>(rabbit<br>monoclonal) | Cell Signaling<br>Technology,<br>Cat #3294 | RRID:AB_823545 | (1:500) |
| Antibody | anti-sXBP-1<br>(rabbit<br>monoclonal) | Cell Signaling<br>Technology,<br>Cat #12782 | RRID:AB_268794<br>3 | (1:500) |
| Antibody | anti-Pax7<br>(mouse<br>monoclonal) | Developmental<br>Studies<br>Hybridoma Bank (<br>DSHB), Cat#<br>pax7, | RRID:AB_528428 | (1:100) |
| Antibody | anti-eMyHC<br>(mouse<br>monoclonal) | DSHB.<br>Cat# F1.652 | RRID:AB_528358 | (1:500) |
| Antibody | anti-MyoD<br>(mouse<br>monoclonal) | SantaCruz<br>Biotechnology,<br>sc-377460 | RRID:AB_281389<br>4 | (1:500) |
| Antibody | anti-Myogenin<br>(mouse<br>monoclonal) | DSHB,<br>Cat# F5D | RRID:AB_528355 | (1:500) |
| Antibody | anti-phospho-p65 (rabbit<br>monoclonal) | Cell Signaling<br>Technology,<br>Cat #3033 | RRID:AB_331284 | (1:500) |
| Antibody | anti-p65<br>(rabbit<br>monoclonal) | Cell Signaling<br>Technology,<br>Cat # 8242 | RRID:AB_108593<br>69 | (1:500) |
| Antibody | anti-p100/p52<br>(rabbit polyclonal) | Cell Signaling Technology,<br>Cat #4882 | RRID:AB_10695537 | (1:500) |
| Antibody | anti-Notch1<br>(mouse monoclonal) | Santa Cruz Biotechnology,<br>Cat # sc-6015 | RRID:AB_831604 | (1:250) |
| Antibody | anti-Hes6<br>(mouse monoclonal) | Santa Cruz Biotechnology,<br>Cat # sc-133196 | RRID:AB_2118109 | (1:250) |
| Antibody | anti-GAPDH<br>(rabbit monoclonal) | Cell Signaling Technology,<br>Cat. #2118 | RRID:AB_561053 | (1:1000) |
| Antibody | anti-laminin<br>(rabbit polyclonal) | Sigma-Aldrich,<br>Cat# L9393 | RRID:AB_477163 | (1:1000) |
| Antibody | anti-Integrin α7-APC (mouse monoclonal) | Miltenyi Biotec,<br>Cat# 130-102-717 | RRID:AB_2652466 | (1:100) |
| Antibody | anti-CD45 Antibody (30-F11), PE,<br>(mouse monoclonal) | ThermoFisher Scientific,<br>Cat # 12-0451-82 | RRID:AB_465668 | (1:400) |
| Antibody | anti-CD31 (PECAM-1) (390), PE,<br>(mouse monoclonal) | ThermoFisher Scientific,<br>Cat # 12-0311-82 | RRID:AB_465632 | (1:400) |
| Antibody | anti-TER-119 (TER-119), PE,<br>(mouse monoclonal) | ThermoFisher Scientific,<br>Cat# 12-5921-82 | RRID:AB_466042 | (1:400) |
| Antibody | anti-Ly-6A/E (Sca-1) (D7), PE,<br>(mouse monoclonal) | ThermoFisher Scientific,<br>Cat # 12-5981-83 | RRID:AB_466087 | (1:400) |

### Animals

C57BL/6 mice and mdx (strain: C57BL/10ScSn-^Dmdmdx^/J) mice were purchased from Jackson Laboratories (Bar Harbor, ME, USA) and breeding colonies were maintained at the University of Houston animal resource facility. Floxed *Ern1* (*Ern1*^fl/fl^) mice as described (Iwawaki et al., 2009) were crossed with *Mck*-Cre mice (Jax strain: B6.FVB(129S4)-Tg (Ckmm-cre)5Khn/J) to generate muscle-specific IRE1α-knockout (i.e. *Ern1*cKO) mice. *Ern1*cKO mice were also crossed with mdx mice to generate littermate mdx;*Ern1*^fl/fl^ and mdx;*Ern1*cKO mice.

Skeletal muscle specific *Xbp1*-knockout mice (*Xbp1*cKO) mice were generated by crossing *Xbp1*^fl/fl^ (MGI strain: *Xbp1*^tm2Glm^) mice as described (Hetz, 2012) with *Mck*-Cre mice. All mice were in the C57BL6 background and their genotype was determined by PCR from tail DNA. We used 10-12 weeks old mice for our experimentation. All the experiments were performed in strict accordance with the recommendations in the Guide for the Care and Use of Laboratory Animals of the National Institutes of Health. All of the animals were handled according to approved institutional animal care and use committee (IACUC) protocols of the University of Houston. All surgeries were performed under anesthesia, and every effort was made to minimize suffering.

### Grip strength measurements

To measure total four-limb grip strength of mice, a digital grip-strength meter (Columbus Instruments, Columbus, OH, USA) was used. In brief, the mice were acclimatized for 5 min and then allowed to grab the metal pull bar with all four paws. The mouse tail was then gently pulled backward in the horizontal plane until it could no longer grasp the bar. The force at the time of release was recorded as the peak tension. Each mouse was tested five times with a 1 min break between tests. The average peak tension from five attempts normalized against total body weight was defined as grip strength.

### Skeletal muscle injury and *in vivo* fusion assay

Muscle necrotic injury in adult mice was performed by injection of 50 µl of 1.2% BaCl_2_ (Sigma Chemical Co.) dissolved in saline into the TA muscle as described (Hindi and Kumar, 2016; Hindi et al., 2012). At various time points after intramuscular injection of BaCl_2_, the mice were euthanized and TA muscle was collected for biochemical and histological studies. To study myoblast fusion *in vivo*, the mice were given an intraperitoneal injection of EdU (4 μg per gram body weight) at day 3 after intramuscular injection of 1.2% BaCl_2_ into the TA muscle. After 11 days of EdU injection, the TA muscle was isolated and sectioned in a microtome cryostat. The sections were subsequently immunostained with anti-Laminin for marking boundaries of myofibers and processed for the detection of EdU^+^ nuclei similar to as described (Hindi et al., 2017). The EdU^+^ nuclei on muscle sections were detected as instructed in the Click-iT^®^ EdU Alexa Fluor^®^ 488 Imaging Kit (Invitrogen). Finally, images were captured and the number of intramyofiber EdU^+^ myonuclei/myofiber, percentage of 2 or more EdU^+^ centrally nucleated fibers and percentage of EdU^+^ myonuclei/total nuclei were quantified using NIH ImageJ software. For calculating the EdU^+^ nuclei, the images were split into single channels and green channel was scanned to count the EdU^+^ nuclei. Similarly, the red and blue channels were scanned to count the numbers of myofibers and myonuclei respectively. While counting, it was ensured that EdU fluorescence is indeed from nucleus. This was achieved by keeping the merged image open in a parallel window and scrutinizing whether the fluorescence overlapped with DAPI and coincided with nuclear location. To reduce variations, three to four different sections from mid-belly of each muscle were included for analysis.

### Histology and morphometric analysis

Uninjured and injured TA muscles were isolated from mice, snap frozen in liquid nitrogen, and sectioned with a microtome cryostat. For the assessment of muscle morphology and to quantify fiber cross-sectional area (CSA), 10μm thick transverse sections of TA muscle were stained with hematoxylin and eosin (H&E). The sections were examined under an Eclipse TE 2000-U microscope (Nikon, Tokyo, Japan). For quantitative analysis, cross-sectional area (CSA) and minimal Feret’s diameter of myofibers was analyzed in H&E-stained TA muscle sections using Nikon NIS Elements BR 3.00 software (Nikon). For each muscle, the distribution of fiber CSA was calculated by analyzing approximately 300 myofibers. Masson’s trichrome staining was performed to analyze fibrosis using a commercially available kit and following a protocol suggested by the manufacturer (Richard-Allan Scientific).

### Isolation and culturing of myofiber

Single myofiber cultures were established from EDL muscle after digestion with collagenase II (Worthington Biochemical Corporation, Lakewood, NJ) and trituration as described (Hindi and Kumar, 2016). Suspended myofibers were cultured in 60 mm horse serum-coated plates in Dulbecco’s modified Eagle’s medium (DMEM) supplemented with 10% fetal bovine serum (FBS; Invitrogen), 2% chicken embryo extract (Accurate Chemical, Westbury, NY), 10 ng/ml basis fibroblast growth factor (Peprotech, Rocky Hill, NJ), and 1% penicillin-streptomycin for three days.

### Immunofluorescence

For the immunohistochemistry studies, frozen TA muscle sections or myofiber were fixed in 4% paraformaldehyde (PFA) in PBS, blocked in 2% bovine serum albumin in PBS for 1h and incubated with anti-Pax7 (1:10, Developmental Studies Hybridoma Bank (DSHB) Iowa City, IA, Cat# pax7, RRID:AB_528428), anti-eMyHC (1:200, DSHB Cat# F1.652 RRID:AB_528358), anti-laminin (1:500, Sigma-Aldrich Cat# L9393 RRID:AB_477163), or anti-MyoD (1:200, Santa Cruz Biotechnology Cat# sc-377460 RRID:AB_631992) in blocking solution at 4°C overnight under humidified conditions. The sections were washed briefly with PBS before incubation with Alexa Fluor 488 (Thermo Fisher Scientific, Cat# A-11034 also A11034 RRID:AB_2576217) or Alexa Fluor 594 (Thermo Fisher Scientific, Cat# A-11037 also A11037 RRID:AB_2534095) secondary antibody for 1 hr at room temperature and then washed 3 times for 5 min with PBS. The slides were mounted using fluorescence medium (Vector Laboratories) and visualized at room temperature on Nikon Eclipse TE 2000-U microscope (Nikon), a digital camera (Nikon Digital Sight DS-Fi1), and Nikon NIS Elements BR 3.00 software (Nikon). Image levels were equally adjusted using Abode Photoshop CS6 software (Adobe).

For quantification of number of Pax7^+^ cells in TA muscle, the images were split into single channels and the red channel was scanned to count the Pax7+ cells. To confirm that red staining is specific to Pax7+ cells, a parallel window with the corresponding merged image was kept open. Care was taken to scrutinize that red fluorescence overlapped with DAPI fluorescence and was also located underneath the basal lamina (laminin staining) to be considered as real Pax7 staining. Similarly, the green and blue channels were scanned to count the numbers of myofibers and nuclei, respectively. For minimizing variation based on muscle size, the satellite cells were quantified from three to four separate regions from the mid-belly of each muscle. A minimum of 500 myofibers were scanned per mouse to count the associated Pax7^+^ cells.

### Fluorescence-activated cell sorting (FACS)

Satellite cells were analyzed by performing FACS analysis as described (Hindi and Kumar, 2016). For satellite cell isolation from heterogeneous cell population, cells were immunostained with antibodies against CD45, CD31, Sca-1, and Ter-119 for negative selection (all PE conjugated, Thermo Fisher Scientific), and with APC-α7- integrin (MBL International) for positive selection..

### Total RNA extraction and qPCR assay

RNA isolation and qPCR were performed using similar protocol as described (Paul et al., 2012; Paul et al., 2010). In brief, total RNA was extracted from uninjured and injured TA muscle of mice using TRIzol reagent (Thermo Fisher Scientific) and an RNeasy Mini Kit (Qiagen, Valencia, CA, USA) according to the manufacturers’ protocols. First-strand cDNA for PCR analyses was made with a commercially available kit (iScript™ cDNA Synthesis Kit, Bio-Rad Laboratories). The quantification of mRNA expression was performed using the SYBR Green dye (Bio-Rad SsoAdvanced™ - Universal SYBR® Green Supermix) method on a sequence detection system (CFX384 Touch Real-Time PCR Detection System - Bio-Rad Laboratories). Primers were designed with Vector NTI software (Thermo Fisher Scientific Life Sciences) and are available from the authors on request. Data normalization was accomplished with the endogenous control (β-actin), and the normalized values were subjected to a 2^-ΔΔCt^ formula to calculate the fold change between control and experimental groups.

### Western blot analysis

Estimation of the presence of various proteins was quantitated by performing Western blot analysis as described (Hindi and Kumar, 2016; Ogura et al., 2015). TA muscle of mice were washed with sterile PBS and homogenized in lysis buffer: 50 mM Tris-Cl (pH 8.0), 200 mM NaCl, 50 mM NaF, 1 mM dithiothreitol, 1 mM sodium orthovanadate, 0.3% IGEPAL, and protease inhibitors. Approximately 100 μg protein was resolved in each lane on 10% SDS-polyacrylamide gels, electrotransferred onto nitrocellulose membranes, and probed with the following antibodies: anti-phospho-IRE1α (1:500; Novus, NB 100-2323), anti-IRE1α (1:500; Cell Signaling Technology, #3294), anti-sXBP-1 (1:1000; Cell Signaling Technology, #12782), anti-Pax7 (1:100; DSHB Cat# pax7, RRID:AB_528428), anti-eMyHC (1:200, DSHB Cat# F1.652 RRID:AB_528358), anti-MyoD (Santa Cruz Biotechnology sc-377460), anti-Myogenin (1:100; DSHB Cat# F5D), anti-phospho-p65 (1:500; Cell Signaling Technology, #3033), anti-p65 (1:500; Cell Signaling Technology, # 8242), anti-p100/p52 (1:500; Cell Signaling Technology, #4882), anti-Notch1 (Santa Cruz Biotechnology, #SC-6015), anti-Hes6 (Santa Cruz Biotechnology, #SC-25396), and anti-GAPDH (1:2000; Cell Signaling Technology, #2118). Antibodies were detected by chemi-luminescence. Quantitative estimation of the bands’ intensity was performed with ImageJ software (NIH).

### Statistical analyses and general experimental design

We calculated sample size using size power analysis methods for a priori determination based on the standard error of mean (SEM) and the effect size was previously obtained using the experimental procedures employed in the study. For animal studies, we estimated sample size from expected number of *Ern1*cKO or *Xbp1*cKO mice and littermate *Ern1*^fl/fl^ or *Xbp1*^fl/fl^ controls. We calculated the sample size for each group as eight animals. Considering a likely drop-off effect of 10%, we set sample size of each group of six mice. For some experiments, three to four animals were found sufficient to obtain statistical differences. Animals with same sex and same age were employed to minimize physiological variability and to reduce SEM from mean. The exclusion criteria for animals were established in consultation with IACUC members and experimental outcomes. In case of death, skin injury, sickness or weight loss of >10%, the animal was excluded from analysis. Muscle tissue samples were not used for analysis in cases such as freeze artefacts on histological section or failure in extraction of RNA or protein of suitable quality and quantity. Animals from different breeding cages were included by random allocation to the different experimental groups. Animal experiments including morphometric analysis of myofiber CSA, percentage of Pax7^+^, or EdU^+^ cells on TA muscle sections were blinded using number codes till the final data analyses were performed. Statistical tests were used as described in the Figure legends. Results are expressed as mean ± SEM. Statistical analyses used two-tailed Student’s t-test or one-way ANOVA followed by Tukey’s multiple comparison test. A value of p<0.05 was considered statistically significant unless otherwise specified.

## DATA AVAILABILITY

All data generated or analyzed during this study are included in the manuscript and supporting files. The source data file with original uncropped western blot images have been uploaded. Source data excel file containing raw data for quantitation graphs has also been uploaded that was used for graphical and statistical analysis in the main manuscript.

## ACKNOWLEDGMENTS

We thank Dr. Laurie Glimcher of Cornell University for providing floxed *Xbp1* mice. This work was supported by funding from NIH grants AR059810 and AR068313 to AK.

## Supplemental Figure Legends

**FIGURE 1-figure supplement 1.**
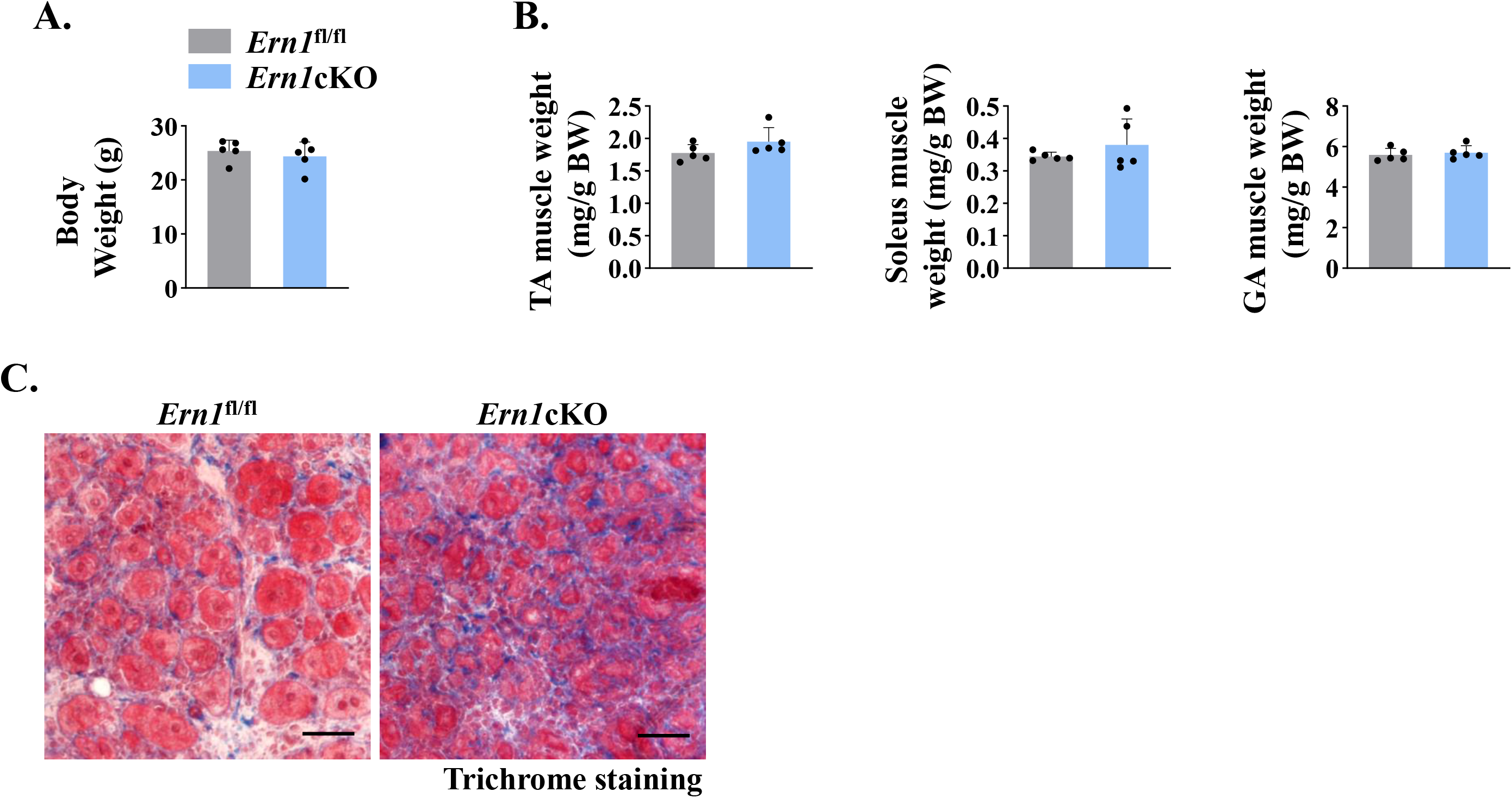
**(A)** Average body weight and **(B)** average wet weight of tibialis anterior (TA
), soleus (Sol), and gastrocnemius (GA) muscle normalized with body weight (BW) of 3-month old *Ern1*^fl/fl^ and *Ern1*cKO mice. **(C)** After 21 days of first injury, TA muscle of *Ern1*^fl/fl^ and *Ern1*cKO mice again given intramuscular injection of 1.2% BaCl_2_ solution, and the muscle was analyzed after 5 days. Representative photomicrograph of TA muscle sections after Masson’s trichrome staining. Scale bar: 50 µm. n=3 mice per group

**FIGURE 4-figure supplement 1.**
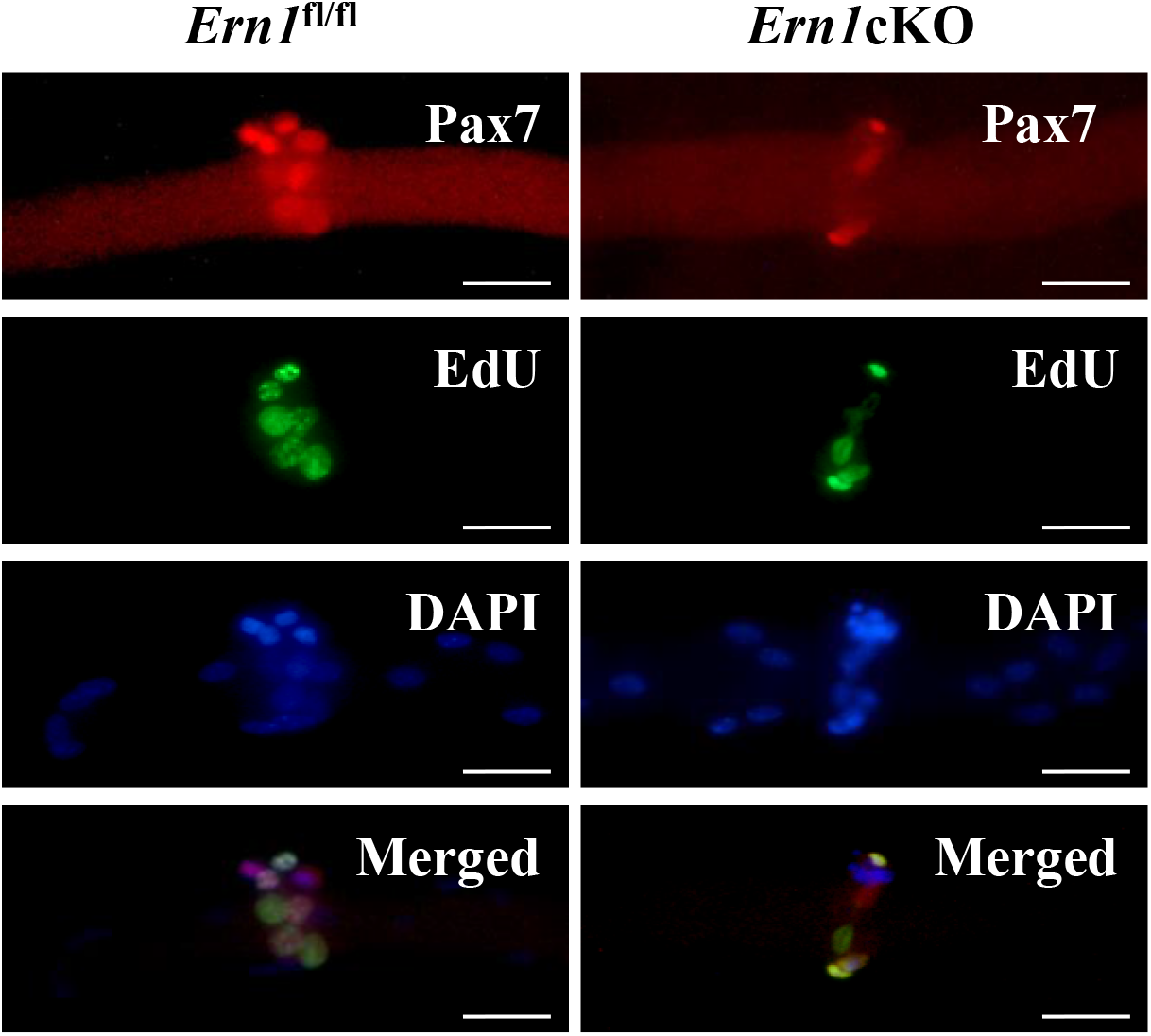
Single myofibers were isolated from the EDL muscle of *Ern1*^fl/fl^ and *Ern1*cKO mice and cultured for 72 h. Following pulse labelling with EdU for 90 min, the myofibers were immunostained for Pax7 protein and EdU. Representative images of myofibers immunostained for Pax7 (red) and EdU (green). Nuclei were counterstained with DAPI (blue). Scale bar: 20 μm.

**FIGURE 5-figure supplement 1.**
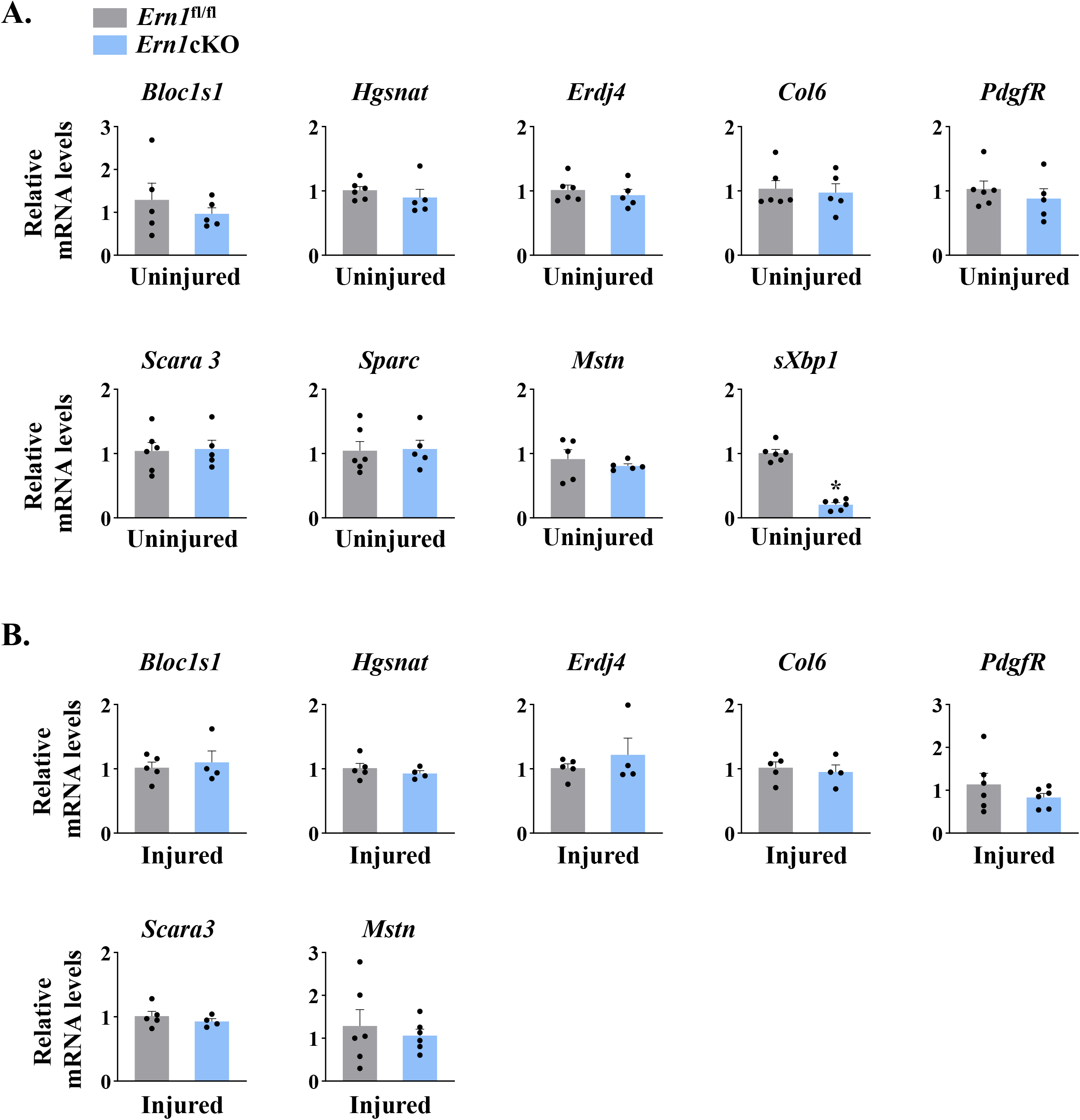
(**A**) Relative mRNA levels of *Bloc1s1*, *Hgsnat*, *Erdj4*, *Col6*, *PdgfR*, *Scara3*, *Sparc*, *Mstn*, and *sXbp1* in uninjured TA muscle of *Ern1*^fl/fl^ and *Ern1*cKO mice (n=5 mice per group). (**B**) Relative mRNA levels of *Bloc1s1*, *Hgsnat*, *Erdj4*, *Col6*, *PdgfR*, *Scara3*, and *Mstn* in 5d-injured TA muscle of *Ern1*^fl/fl^ and *Ern1*cKO mice (n=5 mice for *Ern1*^fl/fl^ group and n=4 mice for *Ern1*cKO group). Data are presented as mean ± SEM. **\***P≤0.05, values significantly different from corresponding uninjured or injured TA muscle of *Ern1*^fl/fl^ mice

**FIGURE 6-figure supplement 1.**
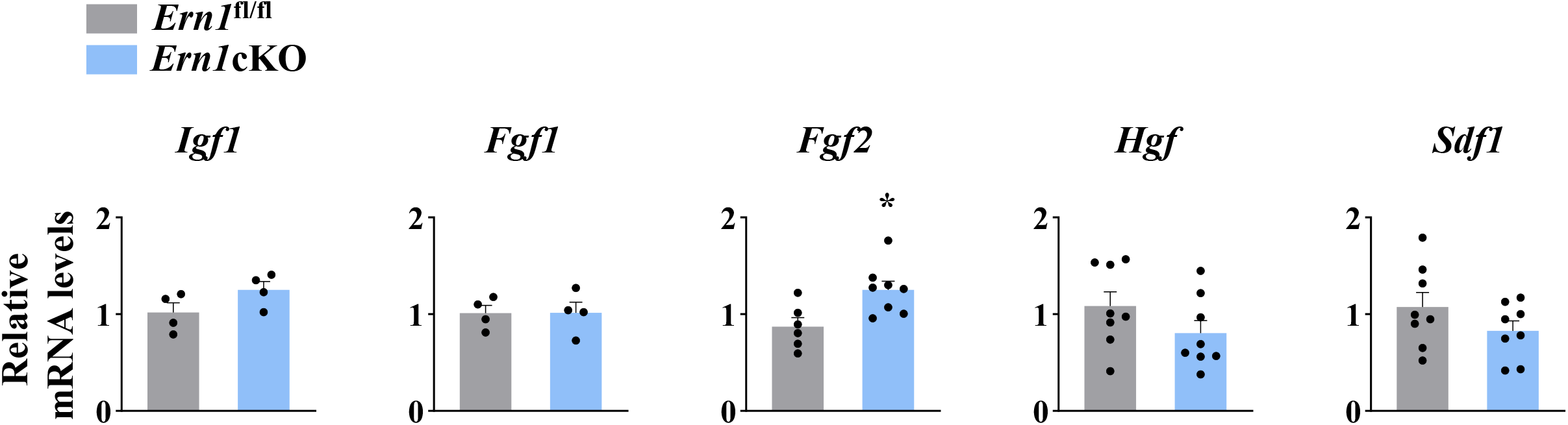
Role of IRE1α in expression of satellite cell growth factors. Relative mRNA levels of *Igf1*, *Fgf1*, *Fgf2*, *Hgf,* and *Sdf1* in 5d-injured TA muscle of *Ern1*^fl/fl^ and *Ern1*cKO mice (n=4-8 mice per group). Data are presented as mean ± SEM. **\***P≤0.05, values significantly different from 5d-injured TA muscle of *Ern1*^fl/fl^ mice.

**FIGURE 8-figure supplement 1.**
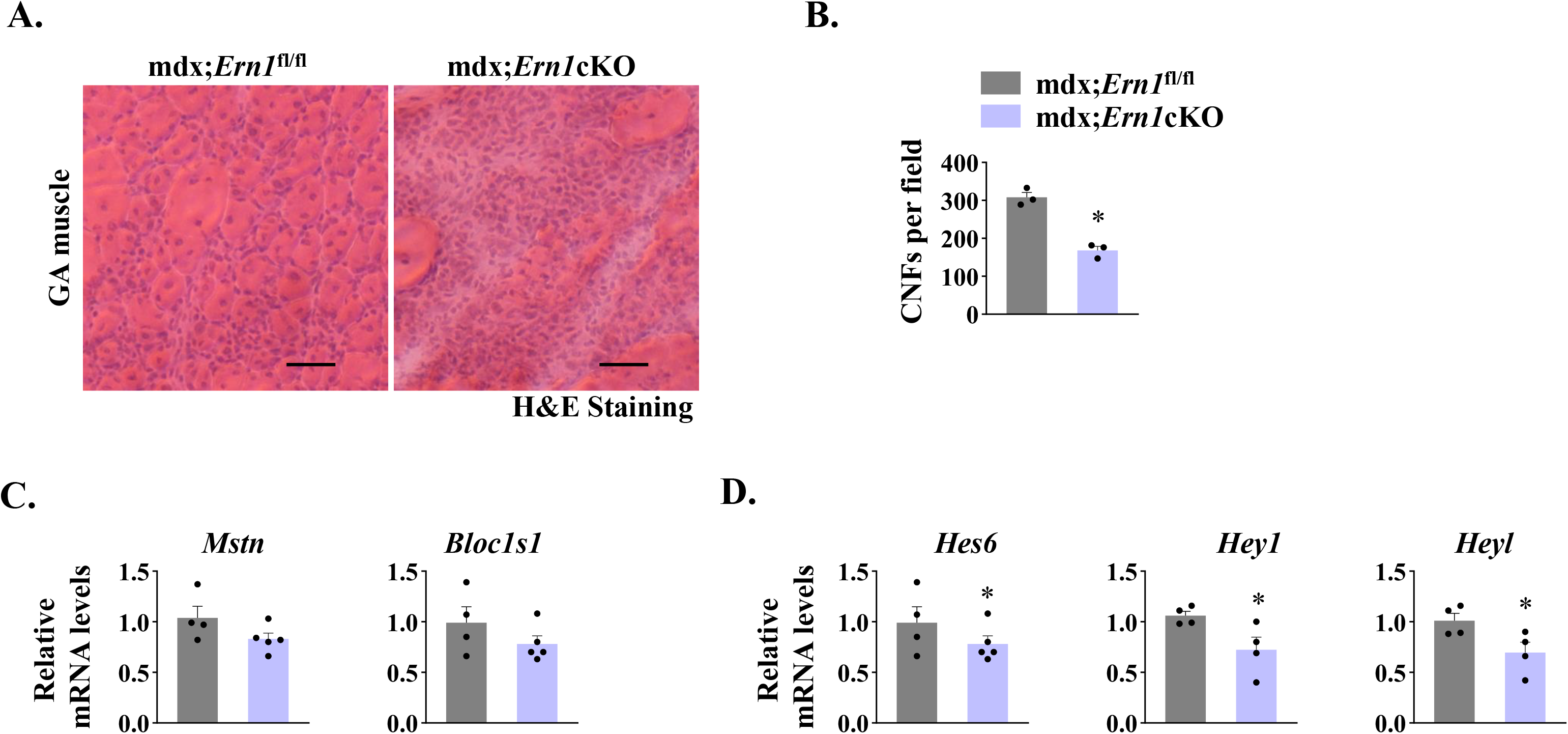
Effect of targeted ablation of IRE1α on muscle pathology in mdx mice. (**A**) Representative photomicrographs of H&E-stained GA muscle sections prepared from 10-month old mdx;*Ern1*^fl/fl^ and mdx;*Ern1*cKO mice. Scale bar: 50 μm. (**B**) Number of centrally nucleated myofibers (CNFs) per field (∼0.15 mm^2^) in GA muscle of mdx;*Ern1*^fl/fl^ and mdx;*Ern1*cKO mice (n=3 mice per group). Relative mRNA levels of (**C**) *Mstn* and *Bloc1s1* (n=4 mice for mdx;*Ern1*^fl/fl^ group and n=5 mice for mdx;*Ern1*cKO group), and (**D**) Notch target genes *Hes6*, *Hey1* and *Heyl* in GA muscle of mdx;*Ern1*^fl/fl^ and mdx;*Ern1*cKO mice (n=4 mice per group). Data are presented as mean ± SEM. **\***P≤0.05, values significantly different from mdx;*Ern1*^fl/fl^ mice by unpaired t test.

